# The redlegged earth mite draft genome provides new insights into pesticide resistance evolution and demography in its invasive Australian range

**DOI:** 10.1101/2022.08.03.502713

**Authors:** Joshua A. Thia, Pasi K. Korhonen, Neil D. Young, Robin B. Gasser, Paul A. Umina, Qiong Yang, Owain Edwards, Tom Walsh, Ary A. Hoffmann

## Abstract

Genomic data provide valuable insights into pest management issues such as resistance evolution, historical patterns of pest invasions and ongoing population dynamics. We assembled the first reference genome for the redlegged earth mite, *Halotydeus destructor* (Tucker, 1925), to investigate adaptation to pesticide pressures and demography in its invasive Australian range using whole-genome pool-seq data from regionally distributed populations. Our reference genome comprises 132 autosomal contigs, with a total length of 48.90 Mb. We observed a large complex of *ace* genes, which has presumably evolved from a long history of organophosphate selection in *H. destructor* and may contribute toward organophosphate resistance through copy number variation, target-site mutations, and structural variants. In the putative ancestral *H. destructor ace* gene, we identified three target-site mutations (G119S, A201S, and F331Y) segregating in organophosphate resistant populations. Additionally, we identified two new *para* sodium channel gene mutations (L925I and F1020Y) that may contribute to pyrethroid resistance. Regional structuring observed in population genomic analyses indicates that gene flow in *H. destructor* does not homogenise populations across large geographic distances. However, our demographic analyses were equivocal on the magnitude of gene flow; the short invasion history of *H. destructor* makes it difficult to distinguish scenarios of complete isolation vs. ongoing migration. Nonetheless, we identified clear signatures of reduced genetic diversity and smaller inferred effective population sizes in eastern vs. western populations, which is consistent with the stepping-stone invasion pathway of this pest in Australia. These new insights will inform development of diagnostic genetic markers of resistance, further investigation into the multifaceted organophosphate resistance mechanism, and predictive modelling of resistance evolution and spread.

## Introduction

The genomes of pest species are providing new evolutionary and applied insights into adaptation to anthropogenic environments. Pesticide resistance in agricultural pests provides an intriguing example of adaptive evolution, where the full genotype-phenotype-environment connection can be made. Toxic pesticides impose selection on populations to evolve phenotypes that can withstand chemical stressors. Because many pesticides exert toxicity through specific biochemical interactions with proteins of target genes, there is a convergence of so called “target-site” mutations across taxa for mutations that block interactions between pesticides and their target gene (Russell *et al*., 2004; Dong *et al*., 2014). Pesticide resistance in agriculture has provided a highly tractable system to study the rise and spread of adaptive variants due to the repeatability of target-site mutations, the ease of characterising adaptive variants, and the large effect size of target-site mutations (Bass *et al*., 2015; Feyereisen *et al*., 2015; Douris *et al*., 2016). Despite their prevalence in agricultural pests, pesticide resistance does not always occur through the evolution of target-site mutations. Metabolic detoxification (Devonshire & Moores, 1982; Walsh *et al*., 2018), behavioural responses (Sparks *et al*., 1989; Zago *et al*., 2014) and reduced penetrance (Gunning *et al*., 1991; Puinean *et al*., 2010) may also contribute to resistance but do not have clear *a priori* genes and (or) alleles. Whilst the genetic basis of behavioural resistance mechanisms is hard to elucidate, the genes underlying metabolic and penetrance resistance mechanisms are more easily identified with access to a reference genome.

In addition to helping resolve the genomic architecture of adaptation, reference genomes support more robust demographic inferences through whole-genome sequencing. Although reduced representation approaches provide efficient ways of obtaining genomic data, their ability to accurately recover the site-frequency spectrum for demographic inference may be dependent on the genomic characteristics of a species, molecular methods implemented and (or) analytical processing (Arnold *et al*., 2013; Rodríguez-Ezpeleta *et al*., 2016; Shafer *et al*., 2016). Reference genomes have aided the investigation of historic demography (Chen *et al*., 2020; Cao *et al*., 2022), invasion routes (Rispe *et al*., 2020; Tay *et al*., 2022) and patterns of admixture (Zhang *et al*., 2020) in agricultural pests globally. From a management perspective, understanding demographic processes operating within pest species can be useful for designing strategies that limit reintroductions from high risk source populations, or for reducing the spread of pesticide resistance (Tay & Gordon, 2019; Thia *et al*., 2021a).

This study focuses on the redlegged earth mite, *Halotydeus destructor* (Tucker 1925; Penthaleidae, Trombidiformes), a significant pest of the grains and pasture industries of Australia. There are two major demographic events in the Australian invasion of *H. destructor*: the first is the original founding event in the state of Western Australia from South Africa around 1917, and the second is the spread of this mite throughout southern regions of Australia during the 1920s (Newman, 1925; reviewed by Ridsdill-Smith, 1997). *Halotydeus destructor* is a winter-active pest, which diapauses as eggs during the Australian summer (October–March), emerges in autumn (April–May), and typically undergoes its active life stage through to late spring (April–November) (Ridsdill-Smith, 1997). Pyrethroid and organophosphate pesticides have been important in the control of *H. destructor* for many decades (Umina *et al*., 2019; Arthur *et al*., 2021). Tolerance to pesticides was first reported in the late 1990s and early 2000s (Hoffmann *et al*., 1997; Robinson & Hoffmann, 2000), with pyrethroid resistance first detected in field populations in 2006 (Umina, 2007) and organophosphate resistance first detected in 2014 (Umina et al. 2017).

Pyrethroid resistance in *H. destructor* has been relatively straightforward to study due to the substantial contribution of a single target-site mutation in the voltage-gated sodium channel gene, *para*, L1024F/V (Edwards *et al*., 2018; Cheng *et al*., 2019; Yang *et al*., 2020). The frequency of this *para* target-site mutation explains 86% to 91% of the variation in pyrethroid resistance among populations of *H. destructor* (Edwards *et al*., 2018). Contrastingly, the study of organophosphate resistance has been challenging. Prior research on *H. destructor* has identified mutations in acetylcholinesterase, *ace*, the target gene of organophosphates (unpublished data). These mutations include G119S, A201S, and F331Y (*Torpedo californica* numbering), which have been observed in a variety of other arthropod species exhibiting target-site resistance. However, complicating this picture have been observations of: (1) susceptible mites carrying these mutations; and (2) evidence of *ace* copy number variation in mite populations, based on quantitative PCR (qPCR) analysis. These observations suggest that organophosphate resistance in *H. destructor* may be conferred by a multifaceted combination of target-site mutations, copy number variation in the *ace* gene, and other additional genomic factors. Follow up investigations have been difficult without a reference genome. Although there have been prior attempts to assemble a reference genome, this assembly was produced from limited short-read data and never fully curated for public use (Edwards *et al*., 2018).

Historically, populations of *H. destructor* have been considered as showing extensive gene flow across Australia, based on allozyme genetic markers (Weeks *et al*., 1995; Qin, 1997). Yet, it is also known that isolation-by-distance patterns can manifest over small geographic distances (<2 km) in this mite, based on reduced-representation genomic data (Yang *et al*., 2020). A long-standing question is whether low levels of genetic differentiation in *H. destructor* are due to high gene flow, large effective population sizes, or a combination of both (Weeks *et al*., 1995). Determining rates of gene flow among populations of *H. destructor* is pertinent for predictive models of resistance evolution dynamics. Whilst there is evidence that resistance often evolves locally (Yang *et al*., 2020) and is constrained by climate (Maino *et al*., 2018), estimates of migration rates across Australia are lacking. Inferring the role of migration in connecting geographically distant populations is an important applied issue with rising pesticide resistance across Australia (Arthur *et al*., 2021).

In this work, we present the first draft genome for *H. destructor*. We used this draft genome to address questions pertinent to resistance management of this mite pest using pool-seq data from regionally distributed populations. Firstly, we examined variation in *ace* copy number and used a comparative analysis to contextualisation observation in *H. destructor* against other arthropod pests. Secondly, we leveraged our genome-wide pool-seq data to screen for target-site mutations in the *ace* and *para* genes. Thirdly, we performed population genomic analyses to characterise the structuring of genetic variation among populations. Fourthly, we performed demographic analyses to assess the role of gene flow in maintaining genetic connections among populations.

## Methods

### *De novo* genome assembly and annotation

#### Biological material and sequencing

To assemble the nuclear genome, samples of adult *H. destructor* were collected from a grassland in Wantirna South, Victoria, Australia (−37°52’11.6", 145°11’46.1”), in June 2020. Collections were via suction using a blower vacuum with a fine gauze mesh placed over the end of the vacuum tube. Mites were placed into a plastic container with paper towel and vegetation for transportation. Once in the laboratory, *H. destructor* individuals were morphologically identified by light microscopy and separated from other invertebrates. DNA was extracted from pooled individuals (*n* = 70) using a DNeasy® Blood & Tissue kit (Qiagen, Hilden, Germany). A continuous long-read library was prepared and sequenced using PacBio Sequel II technology by Berry Genomics (Berry Genomics Co. Ltd, Beijing, China). PacBio sequencing yielded 761,202 reads, with a mean of 13,106 (±6,705 SD) bp, a maximum of 20,938 bp, with ~78% of reads having a length of ≥10,000 bp. Additionally, a NovaSeq 150 bp paired-end Illumina short-read library was prepared and sequenced (Novogene Co. Ltd, Hong Kong), which yielded 20,748,875 reads.

We employed RNA sequence data from a prior study of *H. destructor* (unpublished data) for evidence-based prediction of genes in our draft genome assembly. These transcriptomes came from two populations in Western Australia, one from Capel and one from Manjimup, both of which were collected in 2016 and shown to be resistant to organophosphates through standard pesticide bioassays (Umina *et al*., 2017). Briefly, RNA was extracted using Trizol (Life Sciences, Carlsbad, CA, USA). mRNA was isolated using Oligo d(T)25 Magnetic Beads (New England Biolabs) and fragmented. First strand cDNA synthesis was performed on mRNA fragments with SMARTscribe™ M-MLV Reverse Transcriptase (Takara Bio Inc.). Illumina barcode PCR reactions were performed to prepare the cDNA for Illumina 75 bp paired-end sequencing, which was performed at the Australian National University (Canberra, Australian Capital Territory, Australia). Sequenced transcriptome libraries were quality trimmed with trimmomatic v0.36. In total, 20,183,114 and 22,688,705 paired-end reads were available for the Capel and Manjimup samples, respectively and were *de novo* assembled with trinity v2.2.0 (Grabherr *et al*., 2011).

#### Genome assembly

Raw Illumina reads were analysed with jellyfish v2.3.0 (Marcais & Kingsford, 2012) and genomescope v1.0 (Vurture *et al*., 2017) to perform preliminary genomic profiling in *H. destructor*. Using a kmer size of 21, we inferred a genome size of ~50 Mb, with a good fit between the observed and expected kmer distribution (98%). The heterozygosity rate (the proportion of polymorphic sites relative to total sites) was estimated at 1.4%. Raw PacBio reads were assembled with smartdenovo (Liu *et al*., 2021) to yield a preliminary set of contigs, with a minimum read length threshold of 10,000 bp. We then checked for contamination and polished the genome using an established pipeline (Korhonen *et al*., 2019). For polishing the assembly, Illumina short-reads were quality trimmed using fastp v0.20.1 (Chen *et al*., 2018).

#### Genome annotation

Gene prediction and functional annotation was accomplished using an augustus v3.4.0 software-based pipeline (Korhonen *et al*., 2022), which employs RNA-seq data as evidence data to assist gene prediction. For the annotation of predicted amino acid sequences, InterProScan v5.51-85.0 was used to search the InterPro domain database (Zdobnov & Apweiler, 2001) and blastp v2.9.0 (threshold *E*-value: ≤10−8) was employed to sequence homology databases Swiss-Prot (Magrane, 2011; accessed 30 June 2021), KEGG (Kanehisa *et al*., 2012; accessed 30 June 2021) and NCBI NR (Pruitt *et al*., 2012; accessed 4 February 2021). BUSCO analysis (Simão *et al*., 2015) was used to assess the completeness of our protein annotations. Proteins were screened against the core arthropod (“arthropoda_odb10”) and the arachnid (“arachnida_odb10”) gene sets.

Not all proteins (6,518 sequences) were confidently assigned a GO term. We manually annotated the *ace* copy variants and performed a supplementary GO term assignment with blast2go v5.2.5 (Conesa *et al*., 2005; Conesa & Götz, 2008; Götz *et al*., 2008). All proteins reported for the subclass, Acari, in the NCBI database were downloaded to construct a local BLAST database and mapped against our unannotated genes with blast2go using the default parameters. Collectively, 2,089 additional genes were assigned GO terms. These supplementary annotations are not reported in our NCBI submission but can be found in supplementary Appendix 1.

#### Comparison to other Trombidiformes genomes

To assess the quality of our assembly, we compared our reference genome for *H. destructor* against all available genomes for trombidiform mites, with respect to the number of contigs relative to genome size. We used records of previously assembled genomes using NCBI’s ‘Genome’ search tool, extracting genome assembly stats for all species in the order Trombidiformes (Table 1).

**Table 1.** Previously assembled genomes for mites in the order Trombidiformes.

| GenBank genome | Species | MB | No. contigs | Reference |
| --- | --- | --- | --- | --- |
| GCA_023170135.1 | *Acarapis woodi* | 78.88 | 13685 |  |
| GCA_015350385.1 | *Aculops lycopersici* | 32.54 | 827 | (Greenhalgh *et al.*, 2020) |
| GCA_003956705.1 | *Brevipalpus yothersi* | 71.16 | 2475 | (Denise *et al.*, 2019) |
| GCA_003675995.1 | *Dinotrombium tinctorium* | 180.4 | 25783 | (Dong *et al.*, 2018) |
| GCA_019968535.1 | *Fragariocoptes setiger* | 40.89 | 3583 |  |
| GCA_003675905.2 | *Leptotrombidium deliense* | 117.319 | 66710 | (Dong *et al.*, 2018) |
| GCA_013372475.1 | *Leptotrombidium pallidum* | 170.87 | 190013 | (Kim *et al.*, 2020) |
| GCA_014898815.1 | *Panonychus citri* | 83.97 | 144 | (Yu *et al.*, 2021) |
| GCA_022266195.1 | *Tetranychus cinnabarinus* | 89.72 | 138 | (Huo *et al.*, 2021) |
| == | *Tetranychus urticae* | 85.8 | 3 | (Wybouw *et al.*, 2019) |

### Comparative analysis of *ace* gene sequences

Following annotation, it was apparent that the genome of *H. destructor* harbours many copies of the *ace* gene. We performed a comparative analysis to understand how copy number variation in *ace* varies among other arthropod genomes. *Bemisia tabaci* (Silverleaf whitefly, GenBank GCF_001854935.1; Chen *et al*., 2016), *Leptinotarsa decemlineata* (Colorado potato beetle, GenBank GCF_000500325.1; Schoville *et al*., 2018), *Myzus persicae* (green peach aphid, GenBank GCF_001856785.1; Mathers *et al*., 2017), *Plutella xylostella* (diamondback moth, GenBank GCF_905116875.1) and *T. urticae* (GenBank GCA_000239435.1; Grbić *et al*., 2011) were selected as major agricultural arthropod pests that have evolved resistance to multiple pesticides. *Leptotrombidium deliense* (a species of chigger mite, GenBank GCA_003675905.2; Dong *et al*., 2018) was used as a representative trombidiform mite that is a not an agricultural pest. *Drosophila melanogaster* (GenBank GCA_000001215.4) was also included. Our arthropod gene trees were rooted on the *Torpedo californica ace* sequence (Pacific electric ray, GenBank X03439), which is the canonical reference for the numbering of *ace* target-site mutations.

Downstream analyses were performed in R. First, we constructed a multispecies gene tree to contrast the *H. destructor ace* genes to those in other species. We imported *ace* protein sequences with the packages biostrings v2.62.0 (Pagès *et al*., 2017) and ape v5.6-2 (Paradis *et al*., 2004). Due to truncation and variable length of protein products among *ace* genes, we restricted the multispecies analysis to *ace* genes with proteins ≥500 amino acids in length. ClustalW alignment of *ace* protein sequences was performed with the msa package v1.26.0 (Bodenhofer *et al*., 2015). Following alignment, amino acid sites with missing data were removed, leaving 207 sites for gene tree construction. A distance matrix among protein sequences was derived using the number of amino acid differences. Because we analysed a mix of orthologous and paralogous *ace* sequences in this multispecies analysis, we chose not to implement a model of protein evolution when deriving the distance matrix. We used the ape package to construct a neighbour joining tree from the distance matrix, with node support estimated with 500 bootstrap replicates.

We constructed a second tree to contextualise the *ace* genes in *H. destructor* relative to orthologous versions of the *ace1* and *ace2* genes in other arthropods. We aligned all *H. destructor ace* protein sequences with the *ace1* and *ace2* protein from *B. tabaci* (GenBank EF675187.1 and EF675189.1; Alon *et al*., 2008), *L. decemlineata* (GenBank JF343436.1 and JF343437.1; Revuelta et al., 2011), *M. persicae* (GenBank XM_022304724.1 and AY147797.1). There were no *ace2* orthologs for *L. deliense*, *D. melanogaster*, *P. xylostella*, or *T. urticae* reported in GenBank. Because this alignment included *H. destructor ace* genes with proteins <500 amino acids long, we used pairwise distance to construct the distance matrix for the neighbour joining tree. We used 500 bootstrap replicates to estimate node support.

The R package genomalicious v0.5 (Thia & Riginos, 2019) was used to visualise relative gene positions and structural variation among select genes. The R package *ggtree* v4.2 (Yu *et al*., 2017) was used to visualise trees.

### Target-site mutation screen

We leveraged our reference genome to screen populations for putative resistance conferring target-site mutations at the *ace* and *para* genes. There is a single copy of *para* in the *H. destructor* genome, HDE_05193. However, because of there were many different *ace* genes in the *H. destructor* genome, we focused on *H. destructor ace1* ortholog, HDE_14279 (see Results). We identified non-synonymous SNPs in *para* and *ace* and then assessed whether these mutations had been previously reported to confer resistance in arthropod pests. We used the canonical *Musca domestica para* gene (GenBank NP_001273814.1) and the *T. californica ace* gene (GenBank X03439.1) to reference observed mutations.

### Population genomics

#### Biological material and sequencing

We used our assembled genome to characterise population genetic patterns among Australian *H. destructor* populations. Samples for population genomic analyses came from a study investigating the mechanisms of organophosphate resistance (Table S1). These samples came from populations in the states of Western Australia (Manjimup, Tambellup, Treeton, and Yongarillup) and Victoria (Colbinabbin Res, Colbinabbin Sus; “Res” for resistant, and “Sus” for susceptible to organophosphates). Briefly, DNA from these mites was extracted using a DNeasy® Blood and Tissue Kit (Qiagen, Hilden, Germany) and sequenced using paired-end Illumina 150 bp (Novogene). These data were combined with the pool-seq data from the Wantirna South population used for the genome assembly.

#### Bioinformatics

Reads from each pool-seq library were quality trimmed using fastp and truncated to 100 bp; trimmed reads <100 bp were discarded. After trimming, the mean number of reads per library was 29,570,401 (±3,351,748 SD). Trimmed read pairs were mapped to our draft genome with bowtie2 v2.4.2 using a global alignment and the “--sensitive” mapping option. Samtools v1.12 (Li *et al*., 2009; Danecek *et al*., 2021) was used to discard reads with mapping quality <30 and those that were found to be PCR duplicates. The BAM files from pool-seq libraries from the same population were merged with samtools, and we refer to these merged libraries herein as our “population samples”. In preliminary analyses, we observed considerable variance in depth and coverage of contigs that were <0.2 Mb (Figure S1). Therefore, we limited our reference to those contigs ≥0.2 Mb for population genomic analyses, with a focal set of 56 long contigs.

Variant calling was performed only on those reads mapping to the 56 long contigs. BAM files were subjected to freebayes v1.3.2 (Garrison & Marth, 2012) with the arguments: “—min-alternate-count 3” and “—pooled-continuous”. The freebayes script, fasta_generate_regions.py, was used to segment the genome for variant calling in parallel. We used bcftools v1.12 (Li, 2011; Danecek *et al*., 2021) to combine VCF files from across the parallel variant calling runs. Variant filtering was performed with vcftools v0.1.16 (Danecek *et al*., 2011) to retain only single nucleotide polymorphisms (SNPs; no indels) with a minor allele count of 3, a minor allele frequency of 0.01, minimum quality of 30, minimum depth of 20 reads per sample, no missing data, and to exclude SNPs from repetitive regions of the genome. There were 599,969 SNPs remaining after filtering. The VCF file was imported into R using the package poolfstat v2.0.0 (Hivert *et al*., 2018) and filtered to contain only those SNPs with an overall minor allele frequency of 0.01.

#### Population structure

We calculated the population genomic heterozygosity, *π*, using the estimator 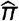 outlined in Ferretti, Ramos-Onsins, & Pérez-Enciso (2013):

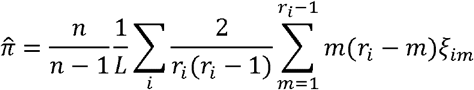

Here, *n* is the number of haploid individuals in a pool, *L* is the length of a focal genomic window, *i* is the index of a focal “cluster” nested within a genomic window, *r_i_* is the read depth within cluster *i*, *m* is the observed alternate read count (ranging from 1 to *r_i_* – 1), and ξ_*im*_ is the number of sites within cluster *i* with *m* counts of the alternate allele. The original formulation of this estimator considers a “cluster” to be a region within a genome with the same read depth, which could be ≥1 bp. For clusters >1 bp, the iteration from *m* = 1 to *r_i_* – 1 for the summation of *m*(*r_i_* – 1)ξ_*im*_ provides a reflection of the site frequency spectrum across multiple adjacent sites with the same read depth. We simplified the calculation by defining clusters within genomic windows as single discrete nucleotide positions (1 bp) that were either polymorphic (a SNP) or monomorphic. This simplifies the calculation because there is a single value of *m* for the *i*^th^ cluster, *m_i_*, and the value of ξ_*im*_ is always 1. And therefore:

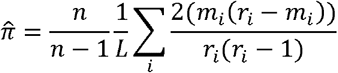

We calculated 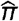 for non-overlapping 5,000 bp windows for our focal set of 56 contigs >0.2 Mb in length. We calculated 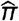 within genomic windows because sequencing depth was not adequate across the entire length of all contigs. We obtained the mean contig 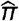 by averaging across windows within each contig, and the mean genome-wide 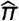 was calculated as the mean of mean contig 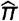 (a “mean of means”). To test the hypothesis that populations varied in their levels of heterozygosity, we constructed a mixed effects model fitting the mean contig 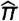 as a function of the fixed “population” effect and the random effect of “contig”. The Wantirna South population was used as the reference level (intercept). Our model was tested using the lme4 v1.1-28 (Bates *et al*., 2014) and lmertest v3.1-3 (Kuznetsova *et al*., 2017) R packages. The multcompview v0.1-8 R package (Graves *et al*., 2019) was used to delineate groups with non-significantly different 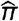 in pairwise *post hoc* comparisons.

Pairwise genome-wide *F*_ST_ was calculated between all population sample combinations using all available SNPs, with the poolfstat package. The poolfstat calculation of *F*_ST_ uses the expected allele frequencies from observed read counts to account for two sources of sampling variance inherent to pool-seq data: estimation of the population allele frequency and estimation of the pooled sample allele frequency. This *F*_ST_ corrected for pool-seq sampling variance is analogous to Weir & Cockerham’s (1984) θ and is calculated as:

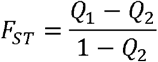

Here, *Q*_1_ is the identity in state probability for two genes sampled within a population pool, and *Q*_2_ is the identity in state probability for two genes sampled between two population pools.

In addition, we used a principal component analysis (PCA) to visualise patterns of population structure. The prcomp function in R was used to conduct a PCA on allele frequency covariances, using the arguments “scale=FALSE, center=TRUE”. We performed two PCAs: one on genome-wide SNPs, and another exclusively on SNPs residing in *ace* genes (exons and introns).

### Demographic inference

To test the importance of gene flow on regional population structure in *H. destructor*, we used forward in time demographic simulations, implemented in the program dadi v2.0.5 (Gutenkunst *et al*., 2009). To model the invasion of *H. destructor* into Australia, we included an ancestral change in population size to represent the bottleneck in the original founding population, a split of the founding population into an eastern and western population, and changes in population size of each population following the east–west split. We then modelled three different scenarios of gene flow: (1) asymmetric migration, where the magnitude of gene flow could vary east-to-west or west-to-east; (2) symmetric migration, where gene flow between regions was equivalent; and (3) isolation, where no gene flow occurred between regions.

Demographic parameters modelled in dadi are relative parameters, scaled by *N*_anc_, the ancestral population size. In our models, the parameter *T*_1_ represented the timing of the ancestral bottleneck in the founding Australian population, with size ν_anc_. The parameter *T*_2_ represented the scaled time of the split of the ancestral population into a western and eastern population, each with size *v*_west_ and *v*_east_. The time parameters were scaled relative to the ancestral population size at the *x*^th^ generation, where *T_x_* = *t_x_*/(2*N*_anc_), with *t_x_* equal to the number of actual generations that have passed. The population size parameters represent the proportional size of the *i*^th^ population relative to the original ancestral population, such that *N_i_* = *v_i_N*_anc_, where *N_i_* is the absolute size of the *i*^th^ population. Finally, gene flow between regional populations was modelled with the scaled migration parameter *M_i,j_*, the number of haploid genomes that move from the *j*^th^ population into the *i*^th^ population per generation. For the isolation scenario, we did not model a migration parameter. For the symmetric migration scenario, gene flow was equivalent in both directions, so we modelled the single parameter, *M*. For the asymmetric migration scenario, we allowed the direction of gene flow to vary by fitting two migration parameters, *M*_east,west_ (west-to-east gene flow) and *M*_west,east_ (east-to-west gene flow). The scaled migration parameters are related to the proportion of migrants per generation, *m_i,j_*, through *M_i,j_* = *m_i,j_N*_anc_.

To infer the values for demographic parameters, we established three eastern versus western population pairs with the following expected haploid genome counts: Wantirna South and Manjimup, *n* =140 and 112; Treeton and Colbinabbin Res, *n* = 84 and 100; and Yongarillup and Colbinabbin Sus, *n* = 164 and 100. We chose Manjimup, Treeton, and Yongarillup to represent western populations because these were genetically more similar to each other then any were to Tambellup (see Results). These working population pairs were made by randomly pairing one eastern with one western population.

Within each population pair, SNPs were filtered for a minor allele frequency of 0.01 and were then thinned to retain one randomly sampled SNP per 5,000 bp. Pooled allele frequencies of thinned SNPs were converted into expected joint site-frequency spectra (SFS) using the dadi_inputs_freqs function from the genomalicious R package, using the argument “methodSFS=‘probs’” to infer allele counts under binomial expectations. Prior to demographic analyses, the joint SFS were downward projected to 20-by-20 haploid genomes with dadi, with final analysed SNP numbers as follows: Wantirna South and Manjimup, 8,368 SNPs; Treeton and Colbinabbin Res, 8,327 SNPs; and Yongarillup and Colbinabbin Sus, 8,313 SNPs. This downward projection was performed to reduce the size of the joint-SFS and reduce computational time.

Demographic models were fit using dadi’s “log_fmin” optimiser, in two phases. The first phase was used to explore parameter space, and the second phase homed in on the most likely parameters. We used ΔAIC criteria to evaluate model support, whereby a pair of models were considered comparable if ΔAIC > −10, and ΔAIC = best model AIC – alternate model AIC (Burnham & Anderson, 2004). We note that this is a conservative threshold for assessing model comparability. We assessed model comparability with ΔAIC at two levels: (1) *within* scenarios, *within* population pairs, to assess the range of parameter values across replicate simulations with similar likelihoods; and (2) *between* scenarios, *within* population pairs, to test support for different demographic scenarios.

In the first phase, we executed 200 replicate simulations, each with 500 maximum iterations, for each scenario and population pair. The seeding parameter values were all set to 0.1. Lower limits for all parameters were set to 1e–10. Upper limits for *T*_1_, *T*_2_, *v*_anc_, *v*_east_, and *v*_west_ were set to 1. Upper limits for *M*, *M*_east,west_, and *M*_west,east_ were set to 10. For each replicate simulation, seeding parameters were perturbed by a factor of 5 within the limits. At the end of the first phase, we identified a set of comparable replicate simulations within scenarios, within population pairs, that had similarly high likelihoods (ΔAIC > −10 relative to the best replicate simulation). The median parameter values across this set of comparable replicate simulations were used to seed parameters in the second phase. In the second phase, we executed 100 replicate simulations, each with 500 maximum iterations. Seeding parameters were perturbed by a factor of 2 and limits were the same as those in the first phase. At the end of the second phase, we again identified the best replicate simulation and those comparable replicate simulations with similarly high likelihoods.

The best replicate simulation from the second phase was used in comparisons between scenarios, within population pairs, as per standard practise. Comparable replicate simulations from the second phase were used to assess the distributions and correlations of parameter values across optimisation runs that achieved similar likelihoods, within scenarios, within population pairs. We note that the primary motivation of our demographic analyses was to try and tease out the importance of migration between eastern and western populations. We did not use these models to infer biologically meaningful parameters, which would require more extensive simulations and genomic bootstrapping (Gutenkunst *et al*., 2009).

## Results

### *De novo* genome assembly and annotation

Our draft *de novo* assembly of the *H. destructor* genome comprised 132 contigs, with a total length of 48.90 Mb, an N50 of 874,492, an L50 of 14, and GC content of 45.3%. The smallest contig was 0.019 Mb and the largest was 4.5 Mb. Note, our assembly does not contain the mitochondrial genome, which we have reported elsewhere (Thia *et al*., 2021b). Our annotation of this genome predicted 14,583 genes. BUSCO analyses indicated that our genome assembly captured a substantial proportion of the core genes expected for arthropods and arachnids. Of the 1,013 single copy orthologs in arthropods, 906 (89.5%) were identified, 892 (88.1%) as single copies and 14 (1.4%) as duplicates; 30 (3.0%) were fragmented and 77 (7.5%) were missing. Of the 2,934 single copy orthologs in arachnids, 2,641 (90%) were found, 2,571 (87.6%) as single copies and 70 (2.4%) as duplicates; 34 (1.2%) were fragmented and 259 (8.8%) were missing. The mapping rate of Illumina short-reads back to genome was 75.58% for the Wantirna South sample that was used to assemble the genome. Our assembly for *H. destructor* was comparable to others obtained for trombidiform mites with respect to number of contigs relative to genome size (Figure 1).

**Figure 1.**
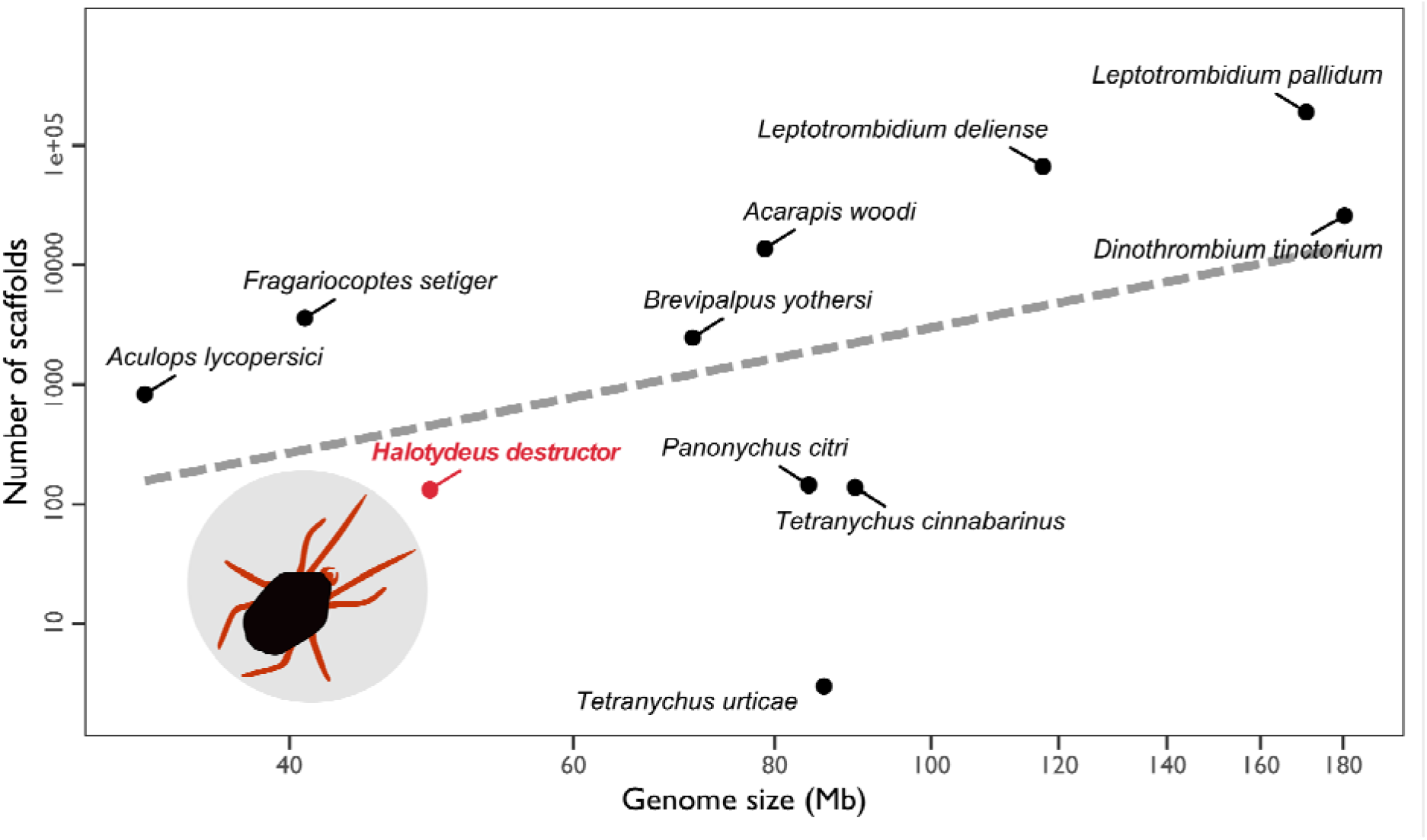
Genome assembly comparison among mites from the order, Trombidiformes. The *x*-axis is the assembled genome size, and the *y*-axis is the number of contigs, both displayed here on a log_10_ scale. The dashed line is the linear fit. Each dark point represents a genome, see Table 1. The red point denotes our assembled *H. destructor* genome. An illustration of *H. destructor* is inset next to its point.

### Comparative analysis of *ace* gene sequences

In total, 28 *ace* genes were identified in the present draft genome for *H. destructor*. The number of *ace* genes in *H. destructor* were markedly higher than that observed in the other arthropod species surveyed, only exceeded by *T. urticae* (46 genes), whereas others had far lower numbers: *B. tabaci* (4 genes), *D. melanogaster* (1), *L. decemlineata* (4), *L. deliense* (10), *M. persicae* (3), and *P. xylostella* (4) (Figure 2). Collectively, the three mites from the order Trombidiformes (*H. destructor*, *L. deliense* and *T. urticae*) had more *ace* genes relative to arthropods in different orders. Most of the *ace* genes in *H. destructor* and *T. urticae* were ≥500 residues in length (25 and 40 *ace* genes), whereas only 2 genes in *L. deliense* were ≥500 residues long (Figure 2). The amplified *ace* genes in *H. destructor* had diverse amino acid sequences, and their divergence from each other was comparable to that observed among species (Figure 3).

**Figure 2.**
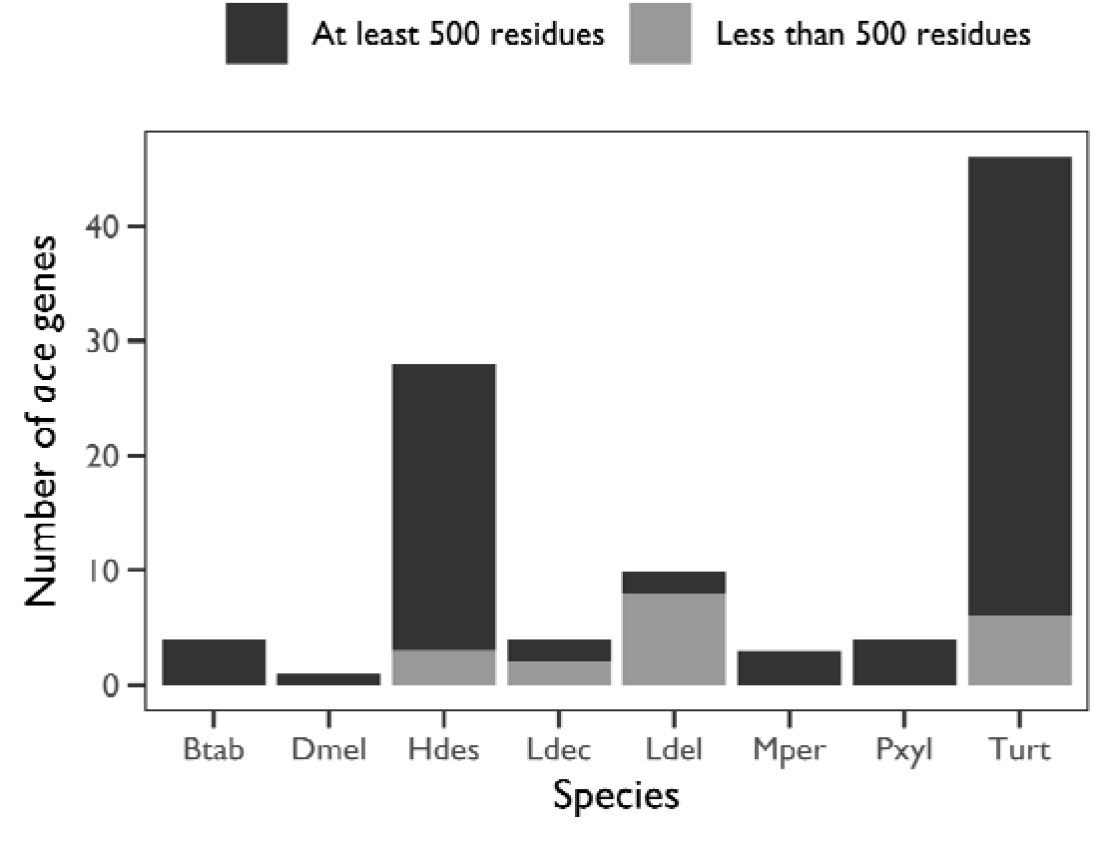
Counts of annotated *ace* genes in select arthropod genomes. Species are on the *x*-axis and the number of annotated *ace* genes in each genome are on the *y*-axis. Bars are coloured by species (see legend). Each panel contains the counts of *ace* genes for different length thresholds: top, all *ace* genes; middle, *ace* genes with a length ≥500 amino acid residues; bottom, *ace* genes with a length <500 amino acid residues. Species abbreviations: “Btab”, *Bemisia tabaci*; “Dmel”, *Drosophila melanogaster*; “Hdes”, *Halotydeus destructor*; “Ldec”, *Leptinotarsa decemlineata*; “Ldel”, *Leptotrombidium deliense*; “Mper”, *Myzus persicae*; “Pxyl”, *Plutella xylostella*; “Turt”, *Tetranycus urticae*.

**Figure 3.**
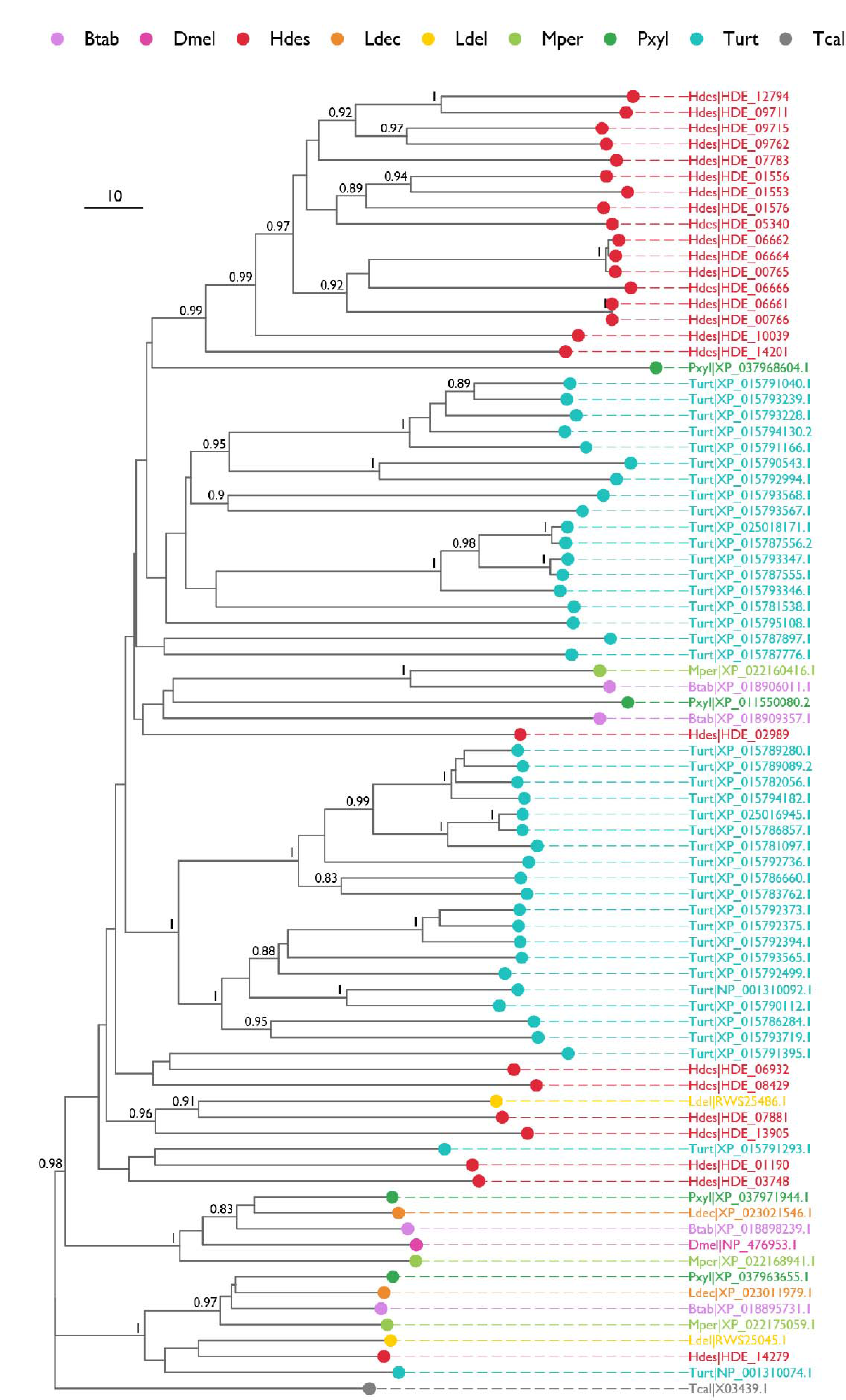
Neighbour joining tree of *ace* protein sequences from the genomes of select arthropod species. Each tip represents an annotated *ace* gene, with points and labels coloured by species (see legend). The *ace* sequence from *Torpedo californica* (“Tcal”) was used to root the tree. Species abbreviations: “Btab”, *Bemisia tabaci*; “Dmel”, *Drosophila melanogaster*; “Hdes”, *Halotydeus destructor*; “Ldec”, *Leptinotarsa decemlineata*; “Ldel”, *Leptotrombidium deliense*; “Mper”, *Myzus persicae*; “Pxyl”, *Plutella xylostella*; “Tcal”, *Torpedo californica*; “Turt”, *Tetranycus urticae*. The scale bar indicates the number of amino acid differences. Nodes labels indicate ≥80% bootstrap support.

The most ancestral *ace* gene in the *H. destructor* genome was HDE_14279, based on its position in the neighbour joining tree relative to other arthropod *ace* genes (Figures 3). Notably, there was a large cluster of 17 *H. destructor ace* genes with shared homology, suggesting a large expansion event from a single gene copy. Such an expansion was also evident in *T. urticae*, with two distinct clusters of *ace* genes present. Homologues of the remaining copies of the *H. destructor ace* genes were present in other species (Figure 3).

There was evidence that some of the *H. destructor ace* gene copies may be associated with structural rearrangements. The clearest example was the *ace* genes found on contig 344 and contig 1156, which exhibit very similar proteins sequences relative to other copies (Figures 3) and appear to be associated with an inversion and deletion event (Figure S2). In our neighbour joining trees of inferred *ace* protein sequences, *ace* genes from contigs 344 and contig 1156 exhibited suspect across-contig relationships (Figure 3). *Ace g*enes HDE_06662 and HDE_06664 on contig 344 were both sister to HDE_00765 on contig 1156, whereas HDE_06661 on contig 344 was sister to HDE_00766 on contig 1156. HDE_06662 and HDE_06664 both differed from HDE_00765 by a total of 66 residues, and HDE_06661 differed from HDE_00766 by just four residues. These groups of related *ace* genes formed a monophyletic group with HDE_06666, which is also on contig 344. Closer examination revealed that contig 1156 (29.53 Kb) comprises a truncated and inverted region of contig 344 (145.58 Kb): contig 1156 starts at the tail-end of contig 344’s HDE_06662 *ace* gene, extending backwards (Figure S2).

However, overall, our assembly likely contains minimal duplication. Both the arthropod (1.4%) and arachnid (2.4%) single copy orthologs in our *H. destructor* assembly were low. We also observed a single copy of elongation factor 1-α (HDE_08018), a common standard for copy number analyses, and a single copy of the *para* sodium channel (HDE_05193), which is the target gene for pyrethroid pesticides. Therefore, we are confident that most of our assembly represents unique genomic regions.

### Target-site mutation screen

We observed two mutations in the *para* gene, and 3 mutations in the ancestral *ace* HDE_14279 gene, that have been reported to confer pesticide resistance in other arthropods (Figure 4). These mutations were only ever found in populations known to be resistant to organophosphates and were absent in both the susceptible Wantirna South and Colbinabbin Sus populations. In the *para* gene, we observed L925I and F1020Y mutations. In the ancestral *ace* gene, we observed G119S, A201S, and F331Y mutations. The mutations ranged in frequency from 0.008 to 0.769.

**Figure 4.**
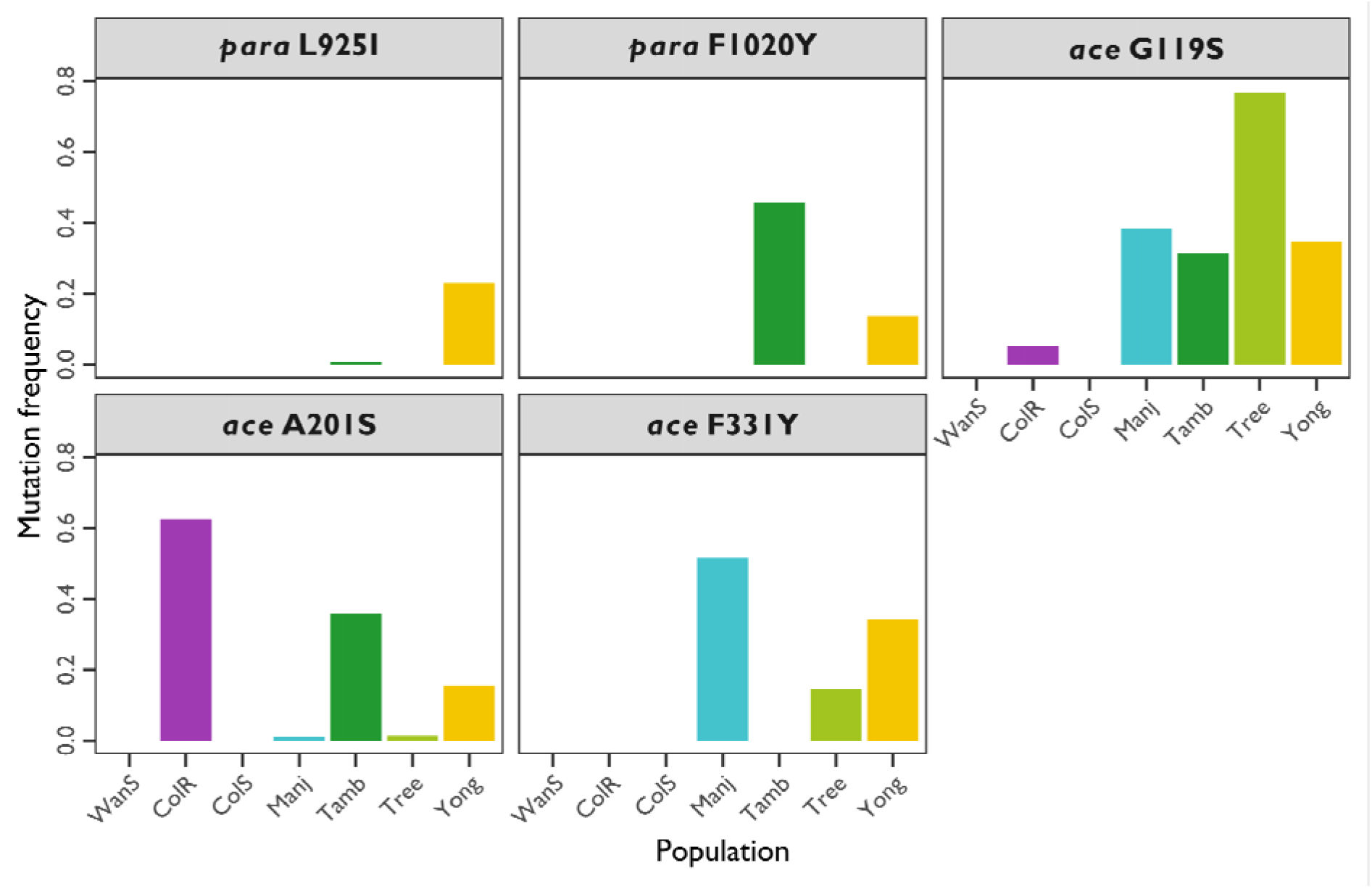
Non-synonymous target-site mutations in the *para* (HDE_05193) and the *ace1* ortholog (HDE_14279) observed in populations of *Halotydeus destructor*. The *x*-axis represents populations, abbreviations as follows: “WanS”, Wantirna South; “ColR”, Colbinabbin Res; “ColS”, Colbinabbin Sus; “Manj”, Manjimup; “Tamb”, Tambellup; “Tree”, Treeton; and “Yong”, Yongarillup. The *y*-axis is the mutation frequency observed in each population. Each panel is mutation.

### Population genomics

We investigated patterns of genome-wide and *ace*-specific population structure within *H. destructor* from the states of Western Australia (western populations) and Victoria (eastern populations) (Figure 5a). The mean genomic heterozygosity, 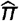, ranged from 0.00416 to 0.00446 (Figure 5b). There were statistically significant differences among populations (χ^2^ = 2,651, df = 7, *p* < 0.001). Overall, eastern populations had lower genome-wide genomic heterozygosity than western populations. Within Victoria, Wantirna South had the greatest diversity 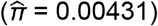, followed by Colbinabbin Res 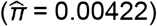, with the least diversity in Colbinabbin Sus 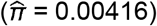. Within Western Australia, genomic heterozygosity was largely comparable among Yongarillup 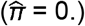, Manjimup 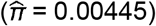, Treeton 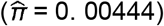, and Tambellup 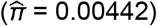. Note, these estimates of genomic heterozygosity reflect the heterozygosity at polymorphic sites standardised by the total number of assayed sites, including monomorphic (Ferretti *et al*., 2013). Such estimates of heterozygosity better reflect genetic diversity and are more robust than SNP-wise heterozygosity that are not standardised for the number of monomorphic sites (Schmidt *et al*., 2021).

**Figure 5.**
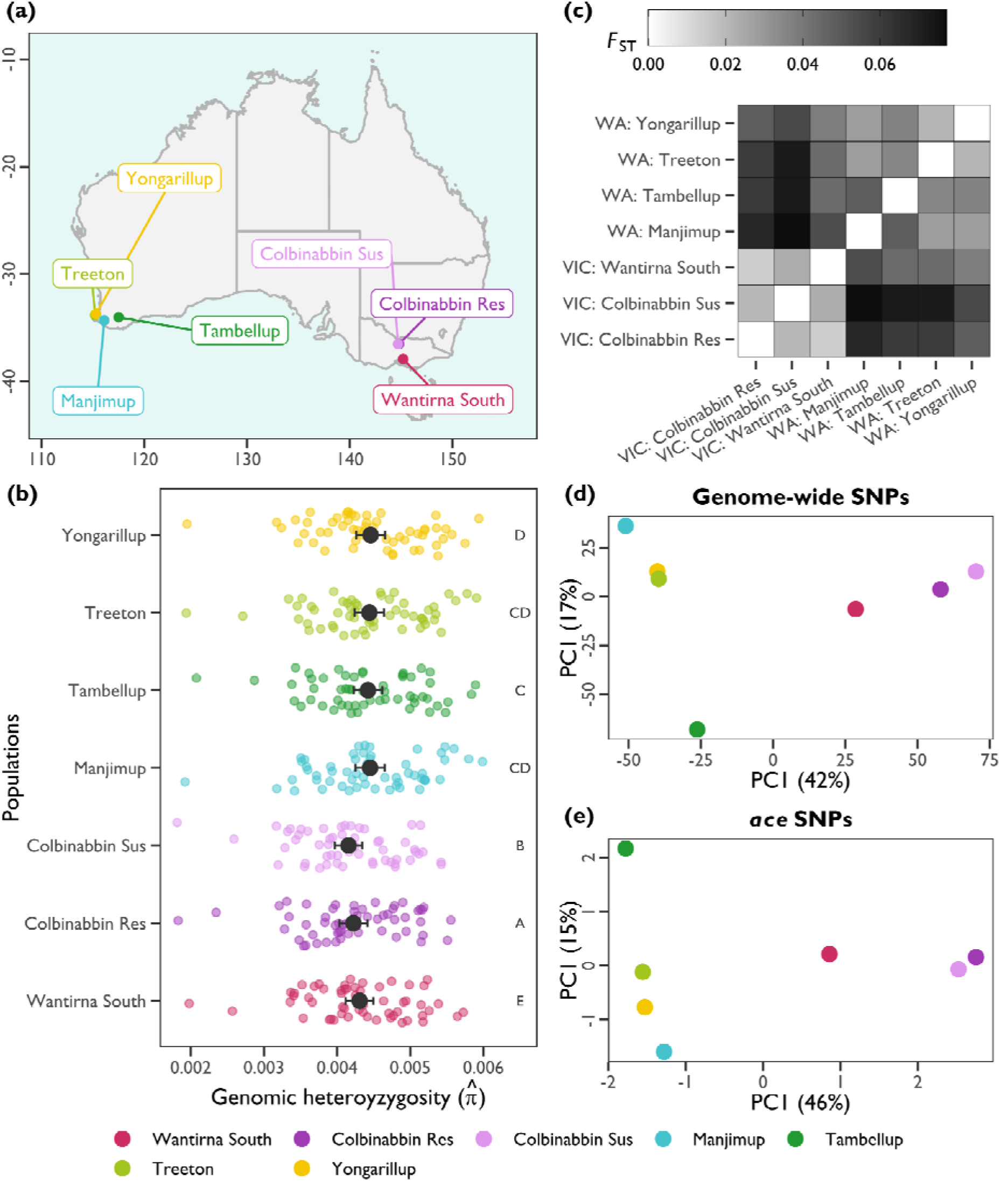
Population genetics of Australian *H. destructor*. (a) Sampling locations of mite populations in Australia: the Manjimup, Treeton, Tembellup, and Yongarillup populations were from state of Western Australia, and the Colbinabbin (Res and Sus) and Wantirna South populations were from the state of Victoria. (b) Population genomic heterozygosity. Populations are on the *x*-axis, with genomic heteroygosity (*π*) on the *y*-axis. Coloured points represent the average *π* for each contig in our draft genome assembly. Large black points represent the average across contigs with 95% confidence intervals. Capital letters indicate groups with non-significant differences in genomic heterozygosity. (c) Pairwise genome-wide *F*_ST_ values among all populations. Darker shading indicates a greater genetic differentiation (see legend). State abbreviations: “VIC”, Victoria; “WA”, Western Australia. (d) Principal component analysis (PCA) of genome-wide allele frequencies. (e) PCA of *ace* gene allele frequencies. (d,e) Points represent a sample, with colour representing the population (see legend). The *x*-axes and *y*-axes represent the first and second principal component, respectively, with numbers in parenthases indicating the percent of explained variance in allele frequencies.

Australian populations of *H. destructor* exhibited a pattern of hierarchical regional population structure (Figure 5c). Pairwise genome-wide *F*_ST_ values ranged from 0.006 014 to 0.0771; they were highest between states (0.038 ≤ *F*_ST_ ≤ 0.077), while they were lower within Western Australia (0.021 ≤ *F*_ST_ ≤ 0.048) and within Victoria (0.014 ≤ *F*_ST_ ≤ 0.023). These data suggest that populations from Western Australia are genetically distinct from those in Victoria, whereas there was little variation among populations within Victoria and some structuring within Western Australia. Within Western Australia, Tambellup was the most differentiated from the other populations. These patterns of differentiation were recapitulated in the PCA of genome-wide SNP allele frequencies (Figure 5d), with regions separated on PC1 and variation within Western Australia separated on PC2. PCA of allele frequencies for *ace* gene SNPs reflected genome-wide patterns, indicating that the local populations contain unique compositions of *ace* variants (Figure 5e).

### Demographic inference

We used the forward in time simulator, dadi, to infer demographic parameters under three modelled scenarios: asymmetric migration, symmetric migration, and isolation (Figure 6a). By our second phase of optimisation, most of the 500 replicate simulations within scenarios and population pairs had reached comparable regions of likelihood space, with ΔAIC > −10 relative to the best model. For the asymmetric migration, symmetric migration, and isolation scenarios (respectively), the number of comparable replicate simulations in phase 2 were as follows: Manjimup and Wantirna South, *n* = 80, 71, and 99; Treeton and Colbinabbin Res, *n* = 57, 71, 99; Yongarillup and Colbinabbin Sus, *n* = 78, 95, and 99 (out of a total of 100).

**Figure 6.**
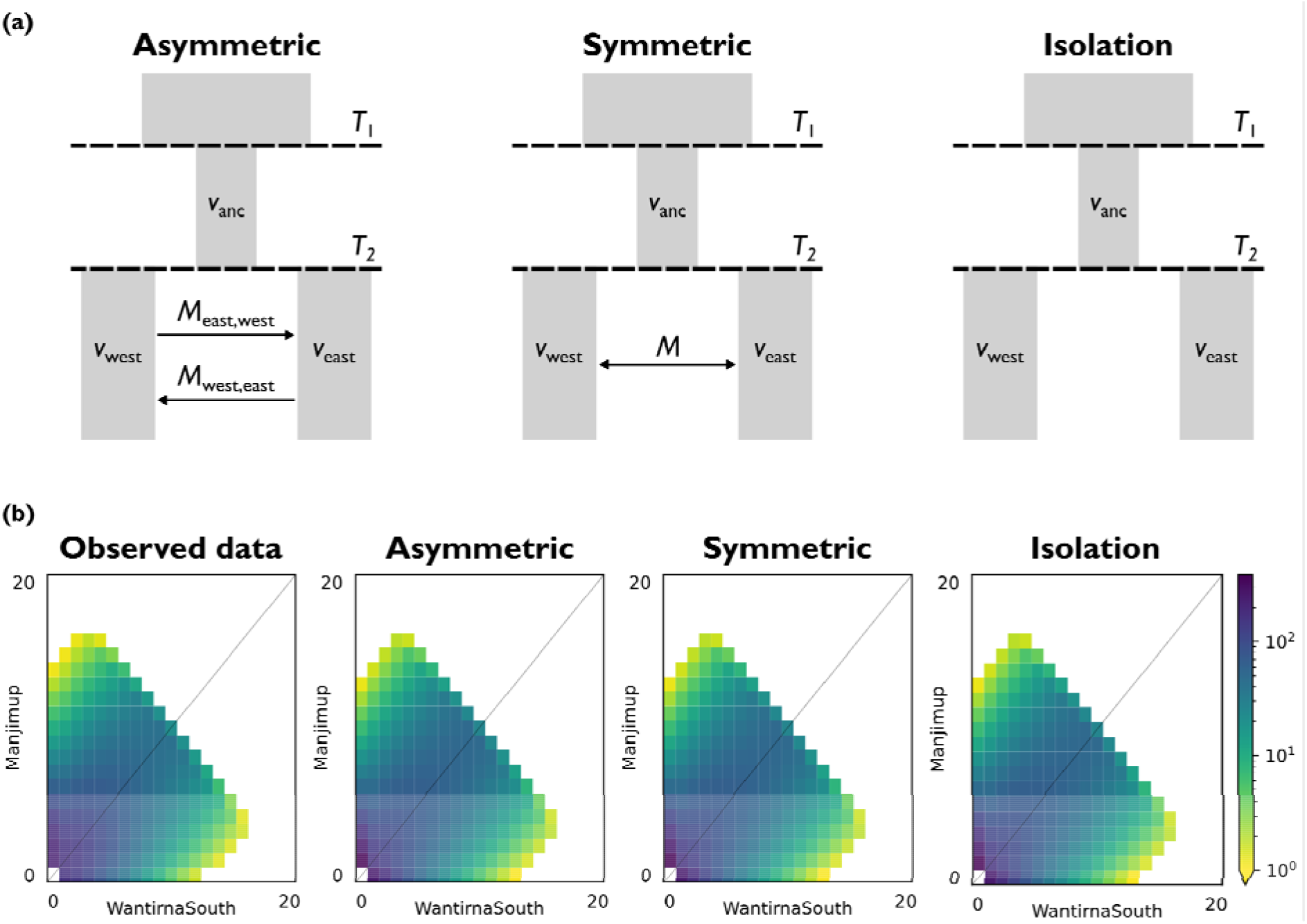
Demographic inference using paired eastern and western populations of *H. destructor*. (a) Schematics of the modelled demographic scenarios and parameters. (b) Representative joint site-frequency spectra (SFS) between Wantirna South (eastern population, *x*-axis) and Manjimup (western population, *y*-axis). Each plot represents the downward projected (20-by-20) joint counts for the minor allele (folded spectra). The scale bar indicates the density of joint allele counts. From left to right, the observed data, and the modelled SFS under the asymmetric migration, symmetric migration, and isolation scenarios. Modelled joint SFS and associated residual plots for all populations pairs can be found in the Supplementary Information (Figures S5–S7).

The best replicate simulations for each scenario and population pair are reported in Table 2. Within population pairs, log-likelihoods of the best replicate simulation were virtually identical for all three scenarios. Across population pairs, and across scenarios, values of *T*_1_, *v*_anc_, *T*_2_, *v*_west_ and *v*_east_ were very small, likely reflecting the recent invasion history of *H. destructor* into Australia and the small sizes of the invasive populations. Western populations had slightly larger inferred population sizes (2.1e–9 ≤ *v*_west_ ≤ 3.6e–8) relative to eastern populations (9.1e–10 ≤ *v*_east_ ≤ 1.8e–8).

**Table 2.** Log-likelihood, AIC, ΔAIC, and demographic parameters for the best replicate simulations for each population pair and scenario. Population abbreviations: ‘Manj’, Manjimup; ‘WanS’, Wantirna South; ‘Tree’, Treeton; ‘ColR’, Colbinabbin Res; ‘Yong’, Yongarillup; ‘ColS’, Colbinabbin Sus.

| Population | Scenario | Log-likelihood | AIC | ΔAIC | $T_1$ | $v_{anc}$ | $T_2$ | $v_{west}$ | $v_{east}$ | $M$ | $M_{west,east}$ | $M_{east,west}$ |
| --- | --- | --- | --- | --- | --- | --- | --- | --- | --- | --- | --- | --- |
| Manj/WanS | Asymmetric | −661.9 | 1337.8 | −4 | 3.5e−10 | 1.7e−09 | 1.0e−10 | 1.9e−09 | 1.1e−09 |  | 1.8e−07 | 1.4e−07 |
|  | Symmetric | −661.9 | 1335.8 | −2 | 1.8e−09 | 8.5e−09 | 1.0e−10 | 1.9e−09 | 1.1e−09 | 5.9e−08 |  |  |
|  | Isolation | −661.9 | 1333.8 | 0 | 1.0e−10 | 4.7e−10 | 1.0e−10 | 1.9e−09 | 1.1e−09 |  |  |  |
| Tree/ColR | Asymmetric | −626.7 | 1267.4 | −4 | 3.7e−09 | 1.8e−08 | 2.0e−09 | 3.6e−08 | 1.8e−08 |  | 3.3e−02 | 3.7E+00 |
|  | Symmetric | −626.7 | 1265.4 | −2 | 1.5e−10 | 7.4e−10 | 1.0e−10 | 1.8e−09 | 9.1e−10 | 1.1e−04 |  |  |
|  | Isolation | −626.7 | 1263.4 | 0 | 1.0e−10 | 4.9e−10 | 1.0e−10 | 1.8e−09 | 9.1e−10 |  |  |  |
| Yong/ColS | Asymmetric | −625.0 | 1263.9 | −4 | 1.0e−10 | 5.1e−10 | 1.0e−10 | 2.1e−09 | 7.5e−10 |  | 8.2e−07 | 5.7e−03 |
|  | Symmetric | −625.0 | 1261.9 | −2 | 1.2e−10 | 6.2e−10 | 1.0e−10 | 2.0e−09 | 7.3e−10 | 3.3e−04 |  |  |
|  | Isolation | −625.0 | 1259.9 | 0 | 1.0e−10 | 5.2e−10 | 1.0e−10 | 2.0e−09 | 7.3e−10 |  |  |  |

Note: See supplementary Figures S6–S11 for distributions and correlations of parameter estimates across comparable replicate simulations.

Although the best supported (lowest AIC) scenario was always the isolation scenario, the symmetric and asymmetric migration scenarios produced comparable fits to the observed SFS (Table 1). In all population pairs, the symmetric migration scenario had ΔAIC = –2, and the asymmetric migration scenario had a ΔAIC = –4, reflecting the cost of adding one and two additional migration parameters, respectively. Indeed, the modelled SFS were indistinguishable among scenarios (Figure 6b) and had identical residual patterns (Figures S3–S5). For the best replicate simulations, estimates of *M*, *M*_west,east_, and *M*_east,west_ were ≤1 (Table 2). However, examination of the parameter estimates across comparable replicate simulations (ΔAIC > −10, relative to best replicate simulation) within each scenario revealed diverse optima under the symmetric and asymmetric migration scenarios. Whilst timing and size parameters exhibited tight convergence across comparable replicate simulations (Figures S6–S8), migration parameters were more diffuse (Figures S6 & S7). Migration rate estimates were not associated with the timing of the ancestral bottleneck, the east–west split, or population sizes (Figures S9 & S10). Furthermore, asymmetric migration rate parameters were uncorrelated; increasing gene flow in one direction was not related to decreasing gene flow in the alternate direction (Figures S9). Our results suggest that inferring the amount and directionality of gene flow in *H. destructor* is challenging because many values of *M*, or *M*_east,west_ and *M*_west,east_ produce comparable fits to the observed SFS within their respective scenarios.

## Discussion

### Draft genome assembly of *Halotydeus destructor*

We report the first draft genome for the redlegged earth mite, *Halotydeus destructor*, a major pest of Australian grains and pastures. This pest is becoming increasingly difficult to manage due to ongoing evolution and spread of pesticide resistance (Arthur *et al*., 2021). Our draft assembly represents the first genome for the mite family Penthaleidae and the eleventh genome for the order Trombidiformes. Genomic resources are under-represented in the Trombidiformes, despite 25,821 described mite species among 151 families (Zhang *et al*., 2011). Our draft genome fills an important gap in available reference genomes. The genome of *H. destructor* is small, only 48.9 Mb in length, which constitutes one of the smallest genomes in the Trombidiformes so far (Figure 1). Of the genomes reported in NCBI, only the gall mite, *Fragariocoptes setiger* (GCA_019968535.1; 40.9 Mb), and the tomato russet mite, *Aculops lycopersici* (GCA_015350385.1; 32.5 Mb), have smaller genomes than *H. destructor*.

Our draft genome was derived from a heterogeneous pool of *H. destructor* from a single location, Wantirna South. Despite the ease of collection, *H. destructor* is challenging to raise under laboratory conditions (Umina, 2007), limiting the production of inbred lines. The increased homozygosity in inbred lines is beneficial for genome assembly, reducing error, fragmentation, and loss of genomic regions (Rhie *et al*., 2021; Whibley *et al*., 2021). Our pooled sample of *H. destructor* used in genome assembly had an estimated heterozygosity rate of 1.4%, the percentage of sites that are polymorphic (*c.f*., genomic heterozygosity, the average genome-wide heterozygosity in a population). Heterozygosity rates in *H. destructor* were comparable to those seen in other trombidiform mites, for example, *Leptotrombidium deliense* (1.2%) and *Dinothrombium tinctorium* (0.7%) (Dong *et al*., 2018). Notwithstanding, our draft genome for *H. destructor* is a relatively complete assembly. For instance, we recovered a major proportion of the expected single copy orthologs for arthropods (89.5%) and arachnids (90.0%). Furthermore, our assembly is on par with other genomes assembled for the Trombidiformes, in terms of the number of scaffolds relative to genome size (Figure 1). Whilst there is no information on the expected chromosome number for mites in the family Penthaleidae, karyotype studies of members of the family Tetranychidae have reported chromosomal numbers between 3 and 14 (Helle & Bolland, 1967; Feiertag-Koppen, 1980; Wybouw *et al*., 2019).

### *Ace* amplification has led to the formation of an *ace* gene complex

Extensive amplification in the acetylecholinesterase gene, *ace*, was revealed by our genome assembly for *H. destructor* (Figures 2 & 3). *Ace* exists as a highly diverse complex of genes. The amino acid differences among *ace* genes in H. *destructor* was comparable to the differences among species (Figure 3). The only surveyed species with more *ace* genes than *H. destructor* was *T. urticae*, one of the most resistant arthropod pests (Van Leeuwen *et al*., 2010; Sparks & Nauen, 2015). It is likely that the *ace* gene complex in *H. destructor* has evolved in responses to decades of organophosphate selection in Australian populations (Arthur *et al*., 2021).

Amplifications of pesticide target genes may be adaptive on two fronts: (1) if increased copy number increases expression, higher pesticide doses are required to suppress the activity of target genes; or (2) by maintaining different susceptible and resistant copies of a target gene, fitness costs associated with target-site mutations can be offset (Feyereisen *et al*., 2015; Weestman *et al*., 2018). In addition to target-site mutations that inhibit interactions between *ace* and organophosphates, *ace* amplifications are also associated with increased resistance in particular strains of *T. urticae* (Khajehali *et al*., 2010; Kwon *et al*., 2010a; b). *Ace* amplifications may also underlie organophosphate resistance in another *Tetranychus* species, *T. evansi*, the tomato red spider mite (Carvalho *et al*., 2012), and are prevalent in organophosphate-resistant mosquito species (Weetman *et al*., 2018). We are currently investigating how copy number variation may influence organophosphate resistance in *H. destructor*.

Not only does the *H. destructor* genome comprise different copies of the *ace* gene, but some of these copies may be associated with structural rearrangements at different genomic scales (Figure S2). Any role of structural variants in pesticide resistance adaptation in *H. destructor* remains to be determined, although chromosomal rearrangements can be important for bringing together genes into adaptive complexes and blocking recombination that might otherwise dissociate these adaptive complexes (Charlesworth & Charlesworth, 1973; Hoffmann *et al*., 2004; Li *et al*., 2022). Understanding the diversity and phenotypic effects of different *ace* structural variants would benefit from long-read sequencing approaches.

### New putative resistance-conferring target-site mutations identified

Our whole-genome sequencing data allowed us unearth candidate target-site mutations for pyrethroid and organophosphate in the *para* and ancestral *ace*, respectively. These two chemicals have been the primary pesticides used for *H. destructor* controls for decades (Ridsdill-Smith *et al*., 2008; Arthur *et al*., 2021). Although pyrethroid resistance via a target-site mechanism evolved some 15 years ago (Umina, 2007), only a single target-site mutation has been reported, the L1024F/V mutation (Edwards *et al*., 2018; Cheng *et al*., 2019; Arthur *et al*., 2021). However, we did not detect this mutation in our study. Instead, we report two new target-site mutations in *H. destructor*, *para* L925I and F1020Y, for the first time. The *para* L925I mutation has been observed in various arthropods (Morin *et al*., 2002; Dang *et al*., 2015; Alissandrakis *et al*., 2017). The F1020Y mutation has not been observed per se, but F1020S mutations have been associated with resistance in diamondback moth (Endersby *et al*., 2011). These *para* target-site mutations were only observed in the Western Australian populations, Tambellup and Yongarillup (Figure 4). However, these populations were not phenotypically screened for pyrethroid resistance, so the role of target-site mutations is currently unclear.

The G119S, A201S, and F331Y *ace* target-site mutations observed in this study have been found in a diversity of organophosphate resistant arthropod pests (e.g., Weill *et al*., 2003; Khajehali *et al*., 2010; Carvalho *et al*., 2012, 2013). These are the first putative target-site mutation reported for organophosphate resistance in *H. destructor*. The frequency of these mutations varied among the resistant populations, and multiple mutations were segregating in the same population (Figure 4). We note that these reported target-site mutations are in the ancestral *ace*, HDE_14279. Although populations appear to harbour unique combinations of *ace* alleles (Figure 5e), the presence of these *ace* HDE_14279 target-site mutations may provide a coarse indicator of resistance risk in *H. destructor*, providing opportunity to develop diagnostic genetic markers (Thia *et al*., 2021a). Nonetheless, allelic variation at other *ace* genes may contribute toward resistance.

### Regional patterns of genetic structure and diversity

Our population genomic analyses revealed genetic differences among populations of *H. destructor* within and between eastern and western regions of Australia (Figure 5c). Among our sampled populations, genetic differentiation was greater between regions (0.038 ≤ pairwise *F*_ST_ ≤ 0.077) relative to within regions (0.014 ≤ pairwise *F*_ST_ ≤ 0.048). Moreover, western populations had significantly higher genomic heterozygosity (Figure 5b) and larger inferred population sizes (Table 2) relative to eastern populations. These observations are consistent with the stepping-stone invasion pathway of *H. destructor* into Australia, whereby the first incursion was in Western Australia, which acted as the source for invasions into eastern regions (Ridsdill-Smith, 1997).

The clear regional structuring of genetic variation suggests that gene flow does not homogenise Australian populations over large geographic distances (Figure 5a & 5c). Quantifying the magnitude of gene flow among populations has been a long-standing question in the management of *H. destructor* (Weeks *et al*., 1995; Qin, 1997). Understanding the rates of dispersal between regions is necessary for predicting the risk of pesticide resistance alleles moving between states. Yang *et al*. (2020) showed that Australian-wide pyrethroid resistance has often evolved through parallel evolution in local populations, but also less frequently via spread of dispersing resistance alleles. This conclusion was based on contrasting genetic differentiation among resistant and susceptible populations at different spatial scales. Yang *et al*. (2020) also found that *H. destructor* can exhibit isolation-by-distance over very short spatial distances (100s of meters), but this does not exclude the possibility of occasional long distance dispersal, such as that required to rapidly spread this pest across Australia within a decade of its introduction (Ridsdill-Smith, 1997).

Our demographic analyses were equivocal on the role of gene flow in shaping patterns of genetic variation. The best model of the joint SFS between eastern and western *H. destructor* populations was always the isolation scenario (Table 2). However, the best replicate simulations for the symmetric and asymmetric migration scenarios had virtually identical fits to the observed data (Figures 6 & S3–S5) with ΔAIC = –2 and –4, respectively. *Halotydeus destructor* invaded Western Australia around 1917, and eastern Australia around 1921 (Ridsdill-Smith, 1997). This short evolutionary history likely poses a challenge for demographic inference; insufficient time makes it difficult to distinguish between scenarios of recent divergence with isolation and ongoing gene flow. Indeed, many values of gene flow produced comparable fits to the observed SFS between eastern and western *H. destructor* populations (Figures S6 & S7). Although a scenario of recent divergence with isolation is the most parsimonious, migration may help maintain some level of genetic connection between regional populations.

The dynamics of *H. destructor* dispersal is expected to be variable; most dispersal is expected to be local and spatially constricted, but also coupled with rare long-distance dispersal events mediated by wind and (or) the movement of livestock, fodder, and machinery. Such a pattern might be validated using individual-level simulations and probabilistic dispersal kernels to compare the effects of ongoing migration with the rare dispersal of propagules between regions. Our ability to infer demographic parameters was limited by our pool-seq design. Whole genome sequences of individuals provide a useful target for future work because they can tap into rich linkage disequilibrium and haplotype information, providing greater power to discern between competing models of gene flow (Sousa & Hey, 2013; Smith & Flaxman, 2020; Marchi *et al*., 2021). Samples from the ancestral South African population would have been useful for contextualising genetic differentiation among invasive Australian populations and inferring demography, but such samples were not available to this study.

### Conclusions

Our draft genome for *H. destructor*, in combination with the recently published mitogenome (Thia *et al*., 2021b), adds to the depauperate genomic resources available for mites in the order Trombidiformes. Our reference genome has provided novel insights into the evolutionary processes operating in the invasive range of *H. destructor*. We have made headway in understanding the multifaceted genomic architecture of organophosphate resistance due to the evolved *ace* gene complex in this mite. Our population genomic investigations have provided greater power to discern subtle regional population differences and refuted the hypothesis that *H. destructor* exists as a single homogeneous population in Australia. Notwithstanding, the nature of the *H. destructor* invasion history will require individual-based data to fully tease out finer details of its demography. Our results will be important in guiding new diagnostic genetic markers for resistance monitoring and for building predictive models of further resistance evolution. We have also generated some exciting new hypotheses on how different *ace* genes may contribute to organophosphate resistance in *H. destructor*.

## Acknowledgements

This work was funded by the Grains Research Development Corporation (Australia) as part of the Australian Grains Pest Innovation Program, and with support from the University of Melbourne. Computational resources were provided by The University of Melbourne’s Research Computing Services and the Petascale Campus Initiative. We thank two anonymous reviewers and editor Mark Ravinet for their constructive feedback.

## Data accessibility

Our draft genome for *H. destructor* is available on GenBank (Assembly, ASM2275052v1; Genome, GCA_022750525.1; Nucleotide, JAIUWE000000000.1). All sequencing data has been submitted under the GenBank Project PRJNA756307. Submissions to the GenBank Sequence Read Archive include: Illumina short-reads for genome assembly (Wantirna South population, SRR15698717); PacBio long-reads for genome assembly (Wantirna South population, SRR15698716), Illumina short-reads for transcriptomes (Capel, SRR20755148; Manjimup, SRR20755147), and Illumina short-reads for population genomics (see Supporting Information, Table S1). All data and scripts associated with these analyses have been deposited into Dryad (Thia, 2022): https://doi.org/10.5061/dryad.xpnvx0kjm.

## Author Contributions

JT, PU, OE, TW and AH contributed to the design of this project. PU and AH obtained funding. JT, OE and PU collected mite populations. OW and TW generated the transcriptome data. JT and QY collected the nuclear genome data. JT, PK, NY and RG contributed to the assembly of the genome. JT led the comparative analyses, population genomics, and demographic modelling. JT wrote the original manuscript. All authors contributed toward revisions of the manuscript.

## Supporting Information

**Figure S1.**
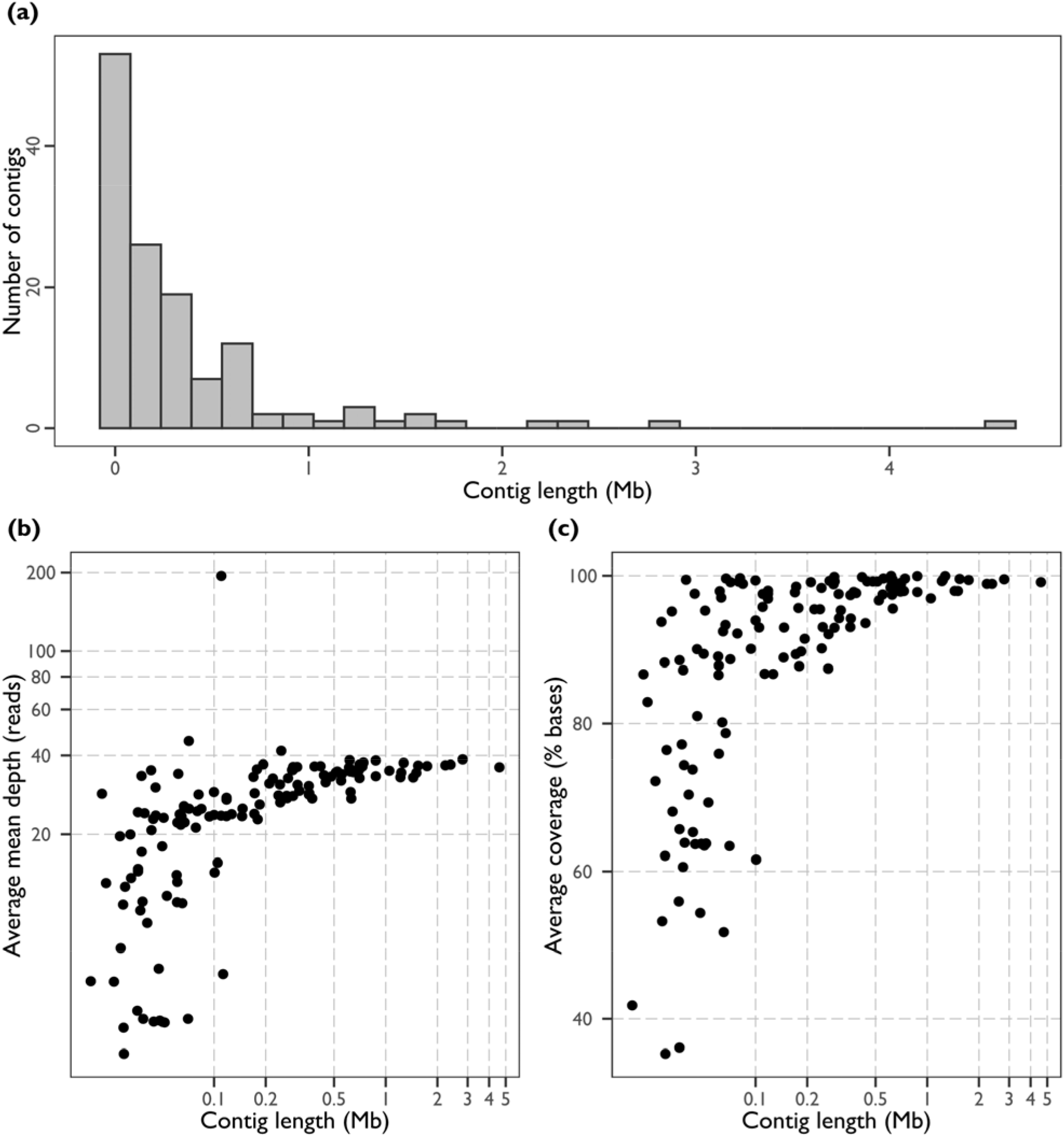
Contig size distributions and mapping efficiency. (a) Distribution of contig sizes in the *Halotydeus destructor* draft genome. (b) Mapping depth across the genome. The *x*-axis is the contig length and the *y*-axis is the average across sample mean depth. That is, for each contig, the mean depth was calculated for each sample, then the average was taken across samples. (c) Average coverage across the genome. The *x*-axis is the contig length and the *y*-axis is the percentage of bases covered.

**Figure S2.**
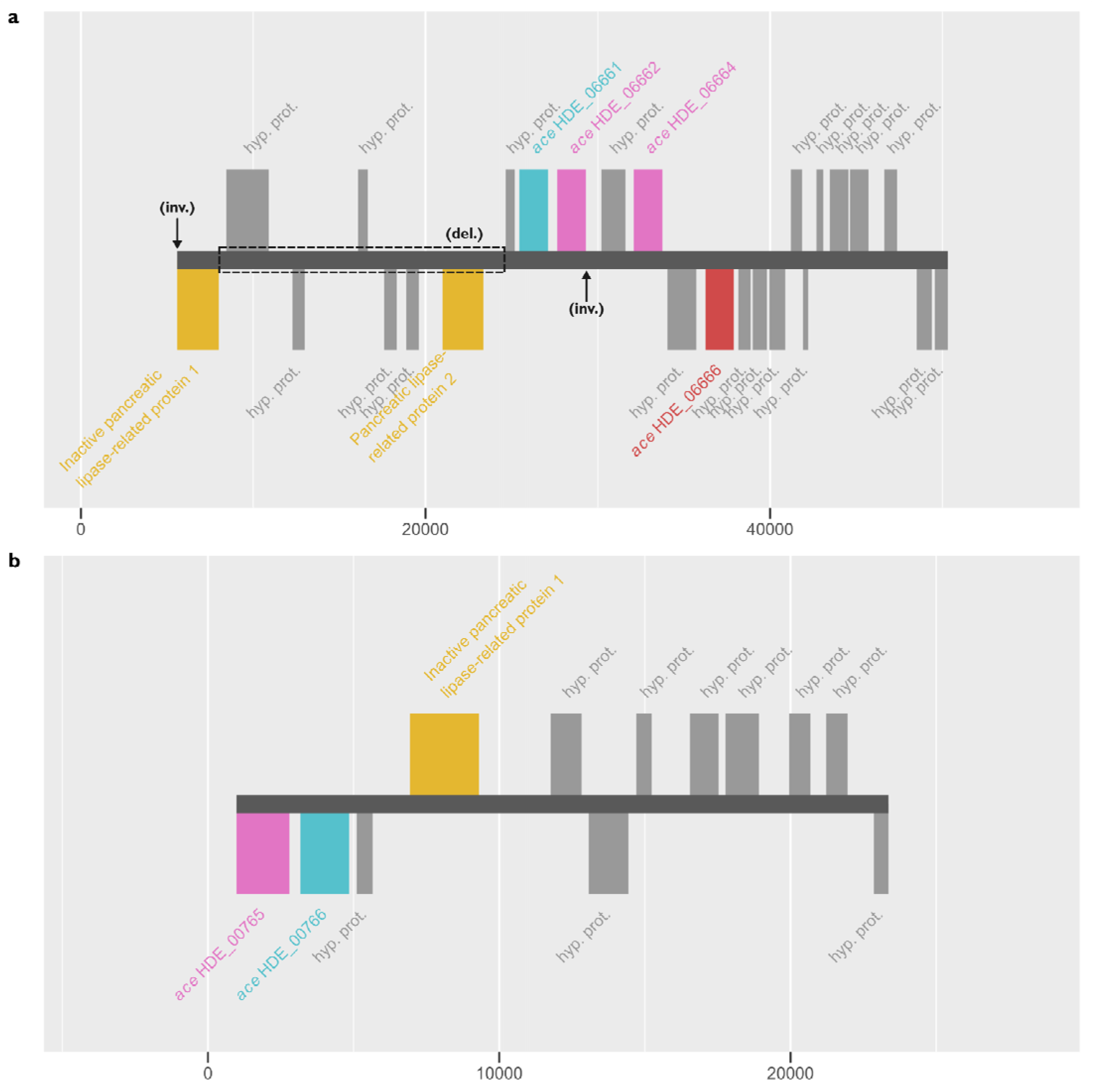
Putative chromosomal inversions of *H. destructor* that contain *ace* genes. The *x*-axis indicates the sequence position (bp) along a genomic contig. The dark grey horizontal bar in the center of the plot represents a subset region of a genomic contig, 20,000 bp either side of focal *ace* genes: (a) contig 344, (b) contig 1156. Blocks situated on the genomic contigs represent annotated genes. Light grey blocks represent hypothetical proteins, “hyp. prot.”. Gold blocks represent two pancreatic lipase-related-like proteins. Blue, pink, and red blocks represent *ace* genes: blue represents the HDE_06661–HDE_00766 sister pair, pink represents the HDE_06662/HDE_06664–HDE_00766 sister pair; and red represents HDE_06666, which is sister to the HDE_06661–HDE_00766 and HDE_06662/HDE_06664–HDE_00766 pairs (see Figure S2). (a) Dashed grey box outlines the region deleted (“del.”) and the arrows point to the ends of the inversion (“inv.”).

**Figure S3.**
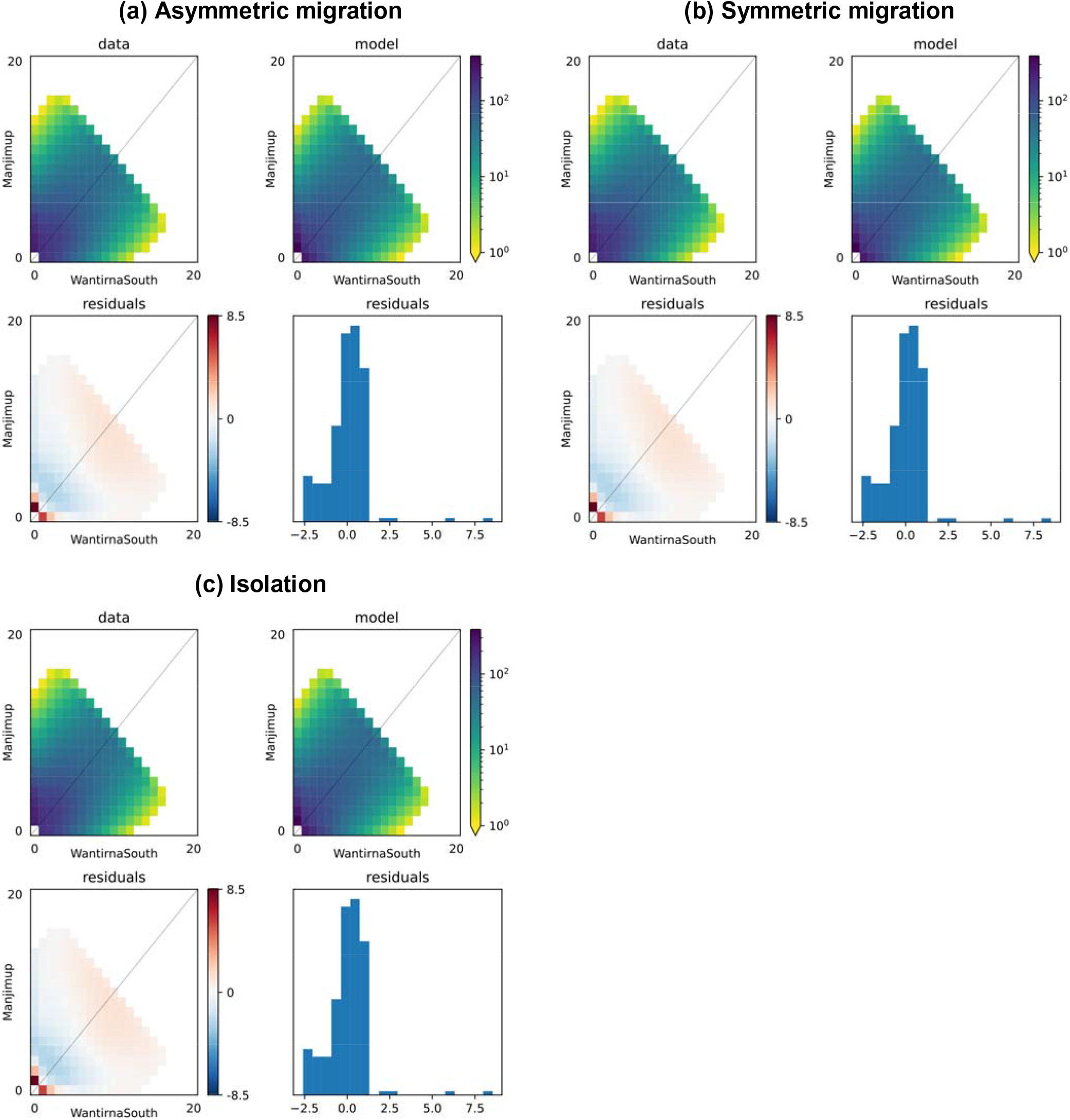
Observed and modelled site frequency spectra (SFS) and residual plots for the best replic simulation, for each modelled scenario, for the Manjimup and Wantirna South population pair. In each subfigure: (top left) The observed SFS; (top right) The modelled SFS; (bottom left) The residuals mapped to the SFS grid; (bottom right) Histogram of residuals. Residuals are the Anscombe residuals, as calculated in *dadi*, effectively depicting the modelled counts minus the observed counts.

**Figure S4.**
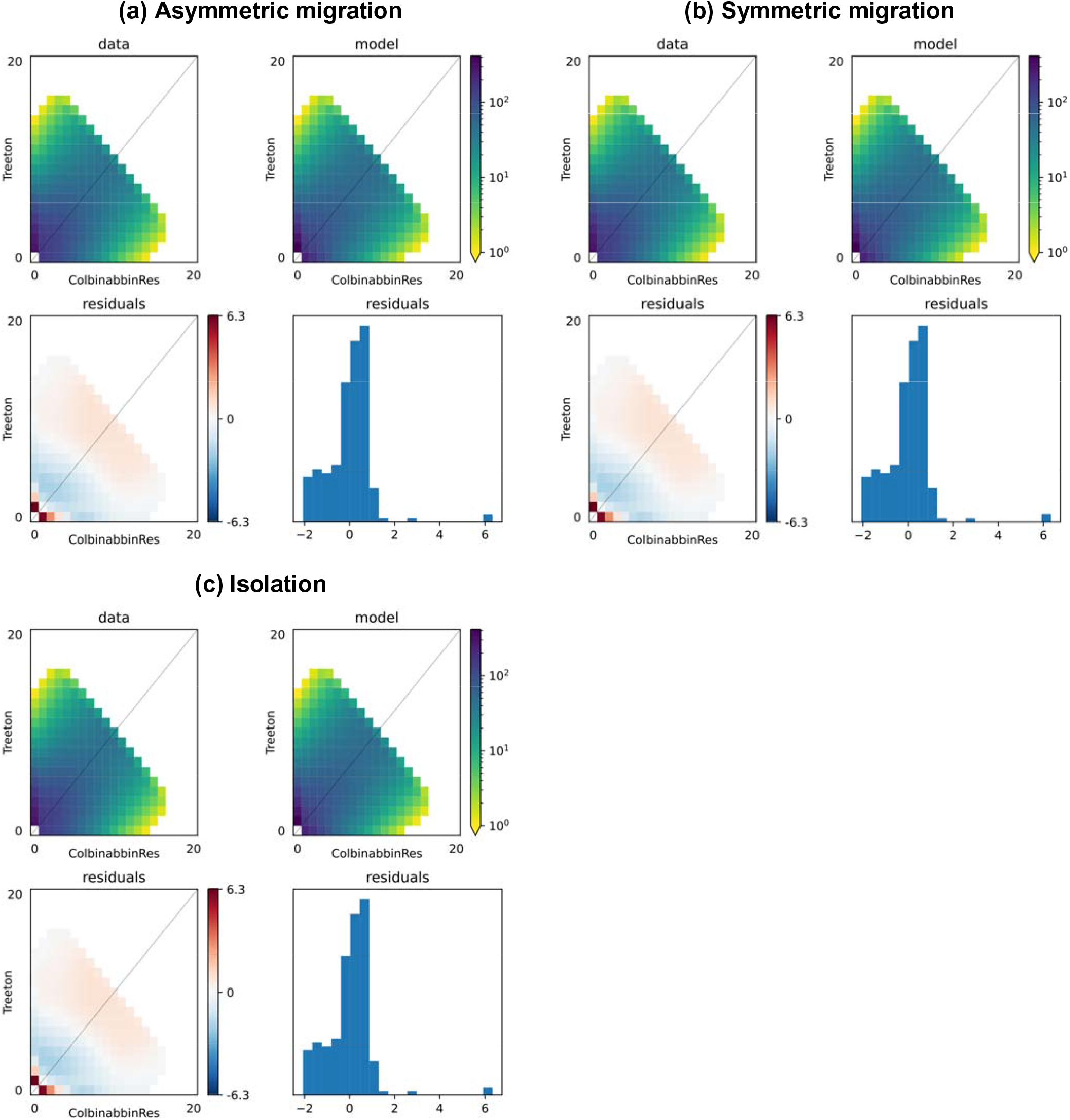
Observed and modelled site frequency spectra (SFS) and residual plots for the best replic simulation, for each modelled scenario, for the Treeton and Colbinabbin Res population pair. In each subfigure: (top left) The observed SFS; (top right) The modelled SFS; (bottom left) The residuals mapped to the SFS grid; (bottom right) Histogram of residuals. Residuals are the Anscombe residuals, as calculated in *dadi*, effectively depicting the modelled counts minus the observed counts.

**Figure S5.**
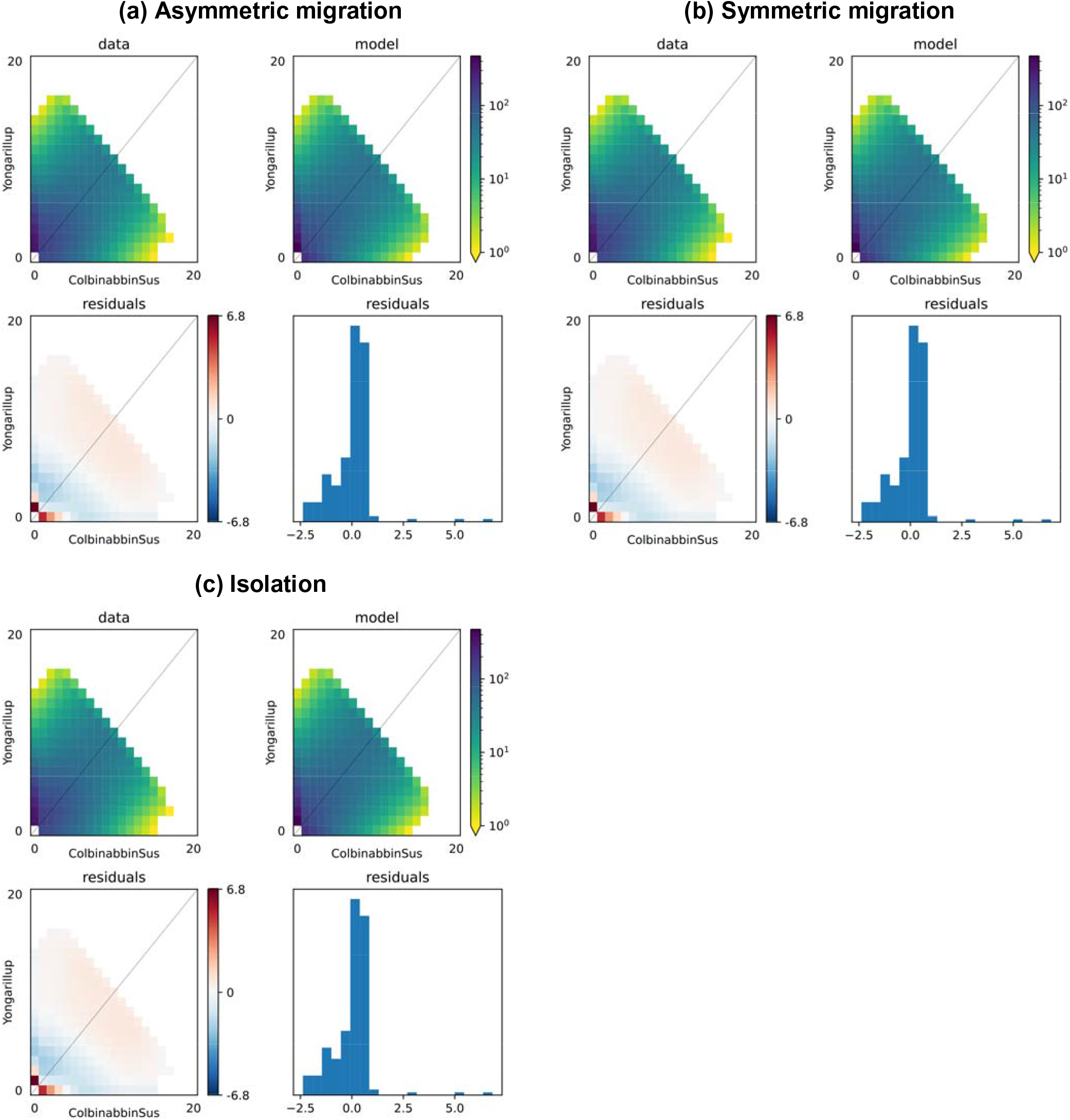
Observed and modelled site frequency spectra (SFS) and residual plots for the best replicate simulation, for each modelled scenario, for the Yongarillup and Colbinabbin Sus population pair. In each subfigure: (top left) The observed SFS; (top right) The modelled SFS; (bottom left) The residuals mapped to the SFS grid; (bottom right) Histogram of residuals. Residuals are the Anscombe residuals, as calculated in *dadi*, effectively depicting the modelled counts minus the observed counts.

**Figure S6.**
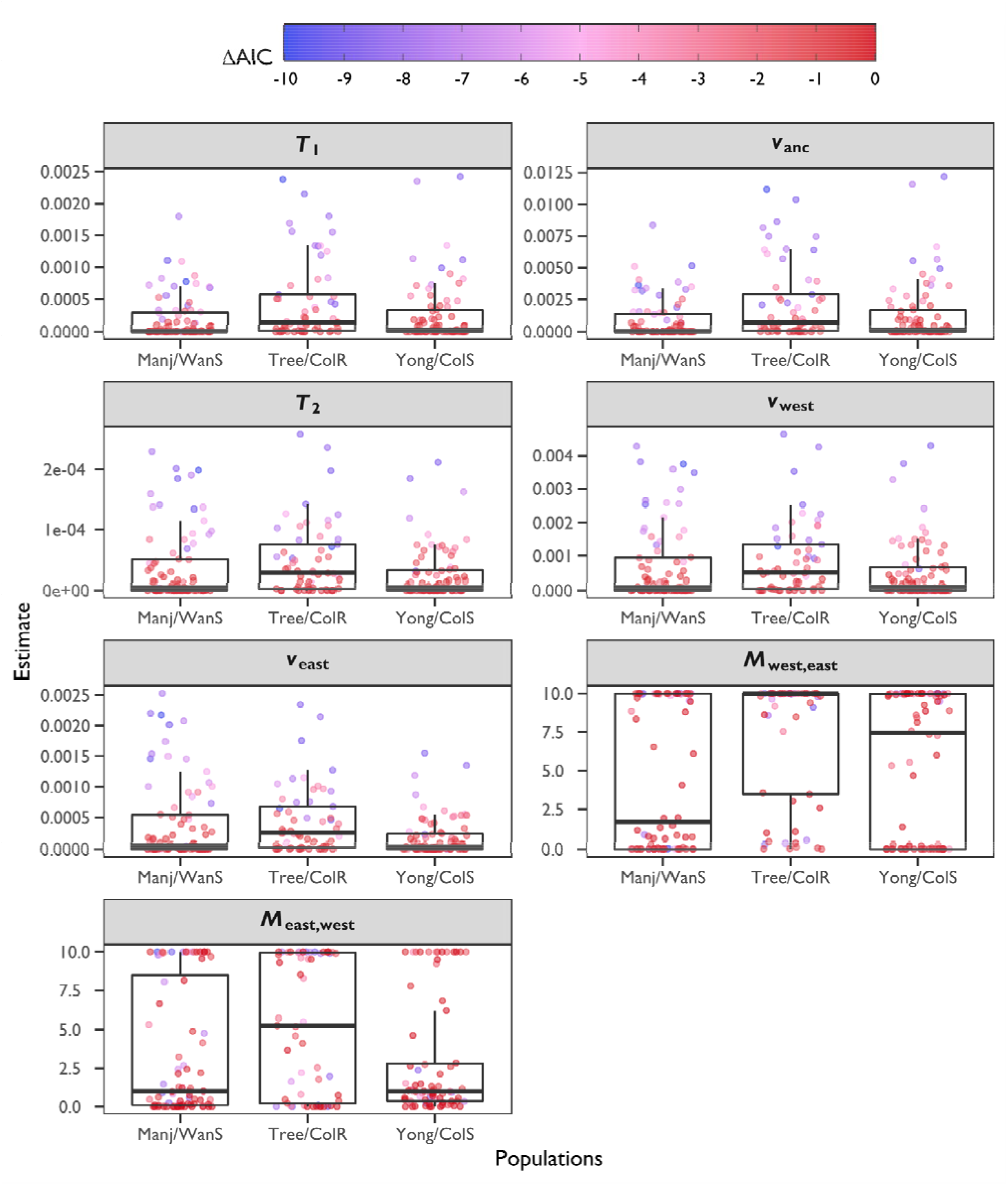
Parameter estimates across comparable replicate simulations (ΔAIC > –10) for the asymmetric migration scenario. Each panel represents a parameter. Population pairs are on the *x*-axis and parameter estimates are on the *y*-axis. Points represent individual replicate simulations, coloured based on their ΔAIC to the best replicate simulation for their population pair (see legend). Boxplots summarise the distribution of estimates. Population abbreviations: ‘Manj’, Manjimup; ‘WanS’, Wantirna South; ‘Tree’, Treeton; ‘ColR’, Colbinabbin Res; ‘Yong’, Yongarillup; ‘ColS’, Colbinabbin Sus.

**Figure S7.**
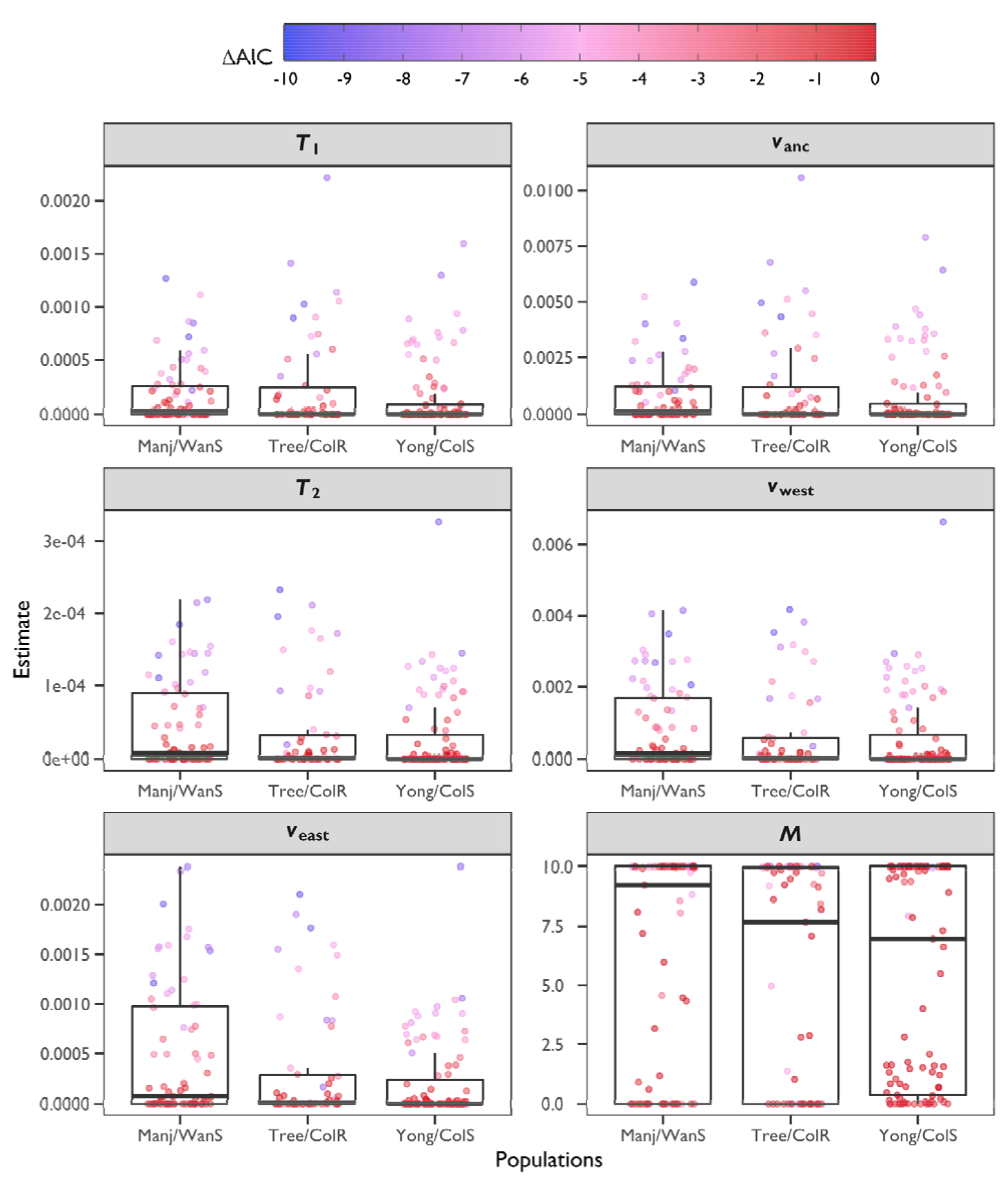
Parameter estimates across comparable replicate simulations (ΔAIC > −10) for the symmetric migration scenario. Each panel represents a parameter. Population pairs are on the *x*-axis and parameter estimates are on the *y*-axis. Points represent individual replicate simulations, coloured based on their ΔAIC to the best replicate simulation for their population pair (see legend). Boxplots summarise the distribution of estimates. Population abbreviations: ‘Manj’, Manjimup; ‘WanS’, Wantirna South; ‘Tree’, Treeton; ‘ColR’, Colbinabbin Res; ‘Yong’, Yongarillup; ‘ColS’, Colbinabbin Sus.

**Figure S8.**
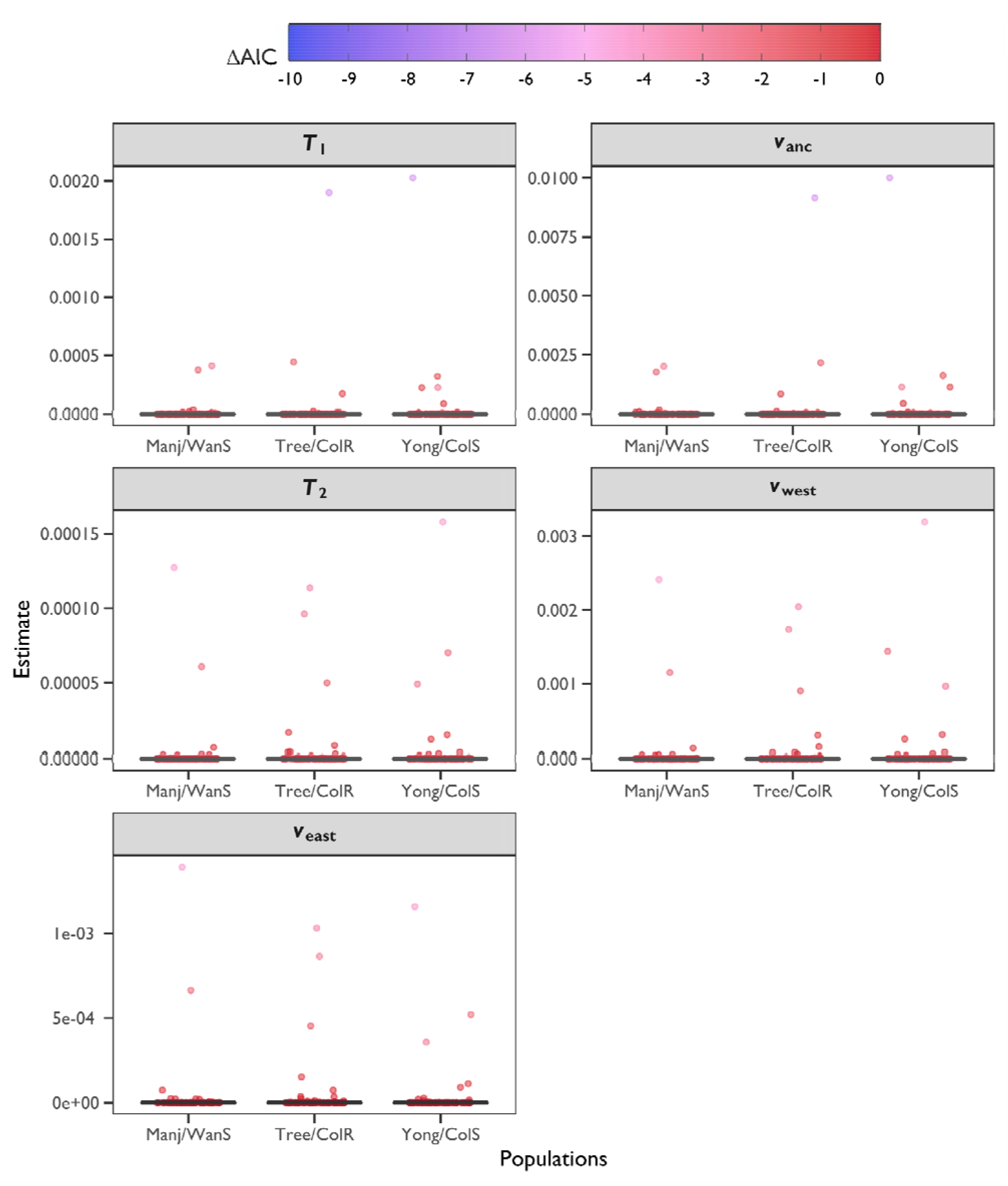
Parameter estimates across comparable replicate simulations (ΔAIC > −10) for the isolation scenario. Each panel represents a parameter. Population pairs are on the *x*-axis and parameter estimates are on the *y*-axis. Points represent individual replicate simulations, coloured based on their ΔAIC to th best replicate simulation for their population pair (see legend). Boxplots summarise the distribution of estimates. Population abbreviations: ‘Manj’, Manjimup; ‘WanS’, Wantirna South; ‘Tree’, Treeton; ‘ColR’, Colbinabbin Res; ‘Yong’, Yongarillup; ‘ColS’, Colbinabbin Sus.

**Figure S9.**
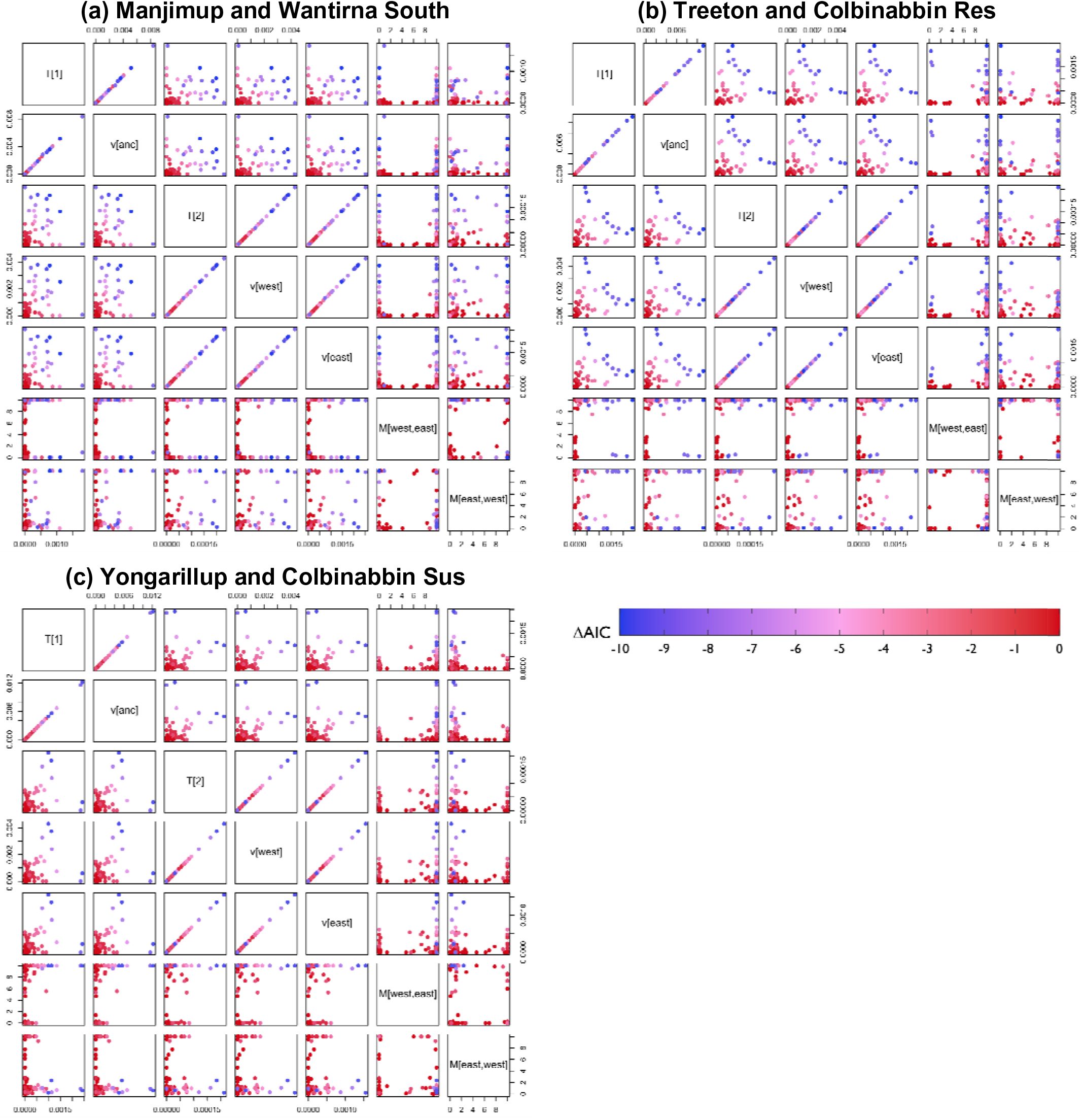
Correlation among parameter estimates for comparable replicate simulations (ΔAIC > −10) modelled under the asymmetric migration scenario, for each population pair. Points represent individual replicate simulations, coloured based on their ΔAIC to the best replicate simulation for their population pair (see legend).

**Figure S10.**
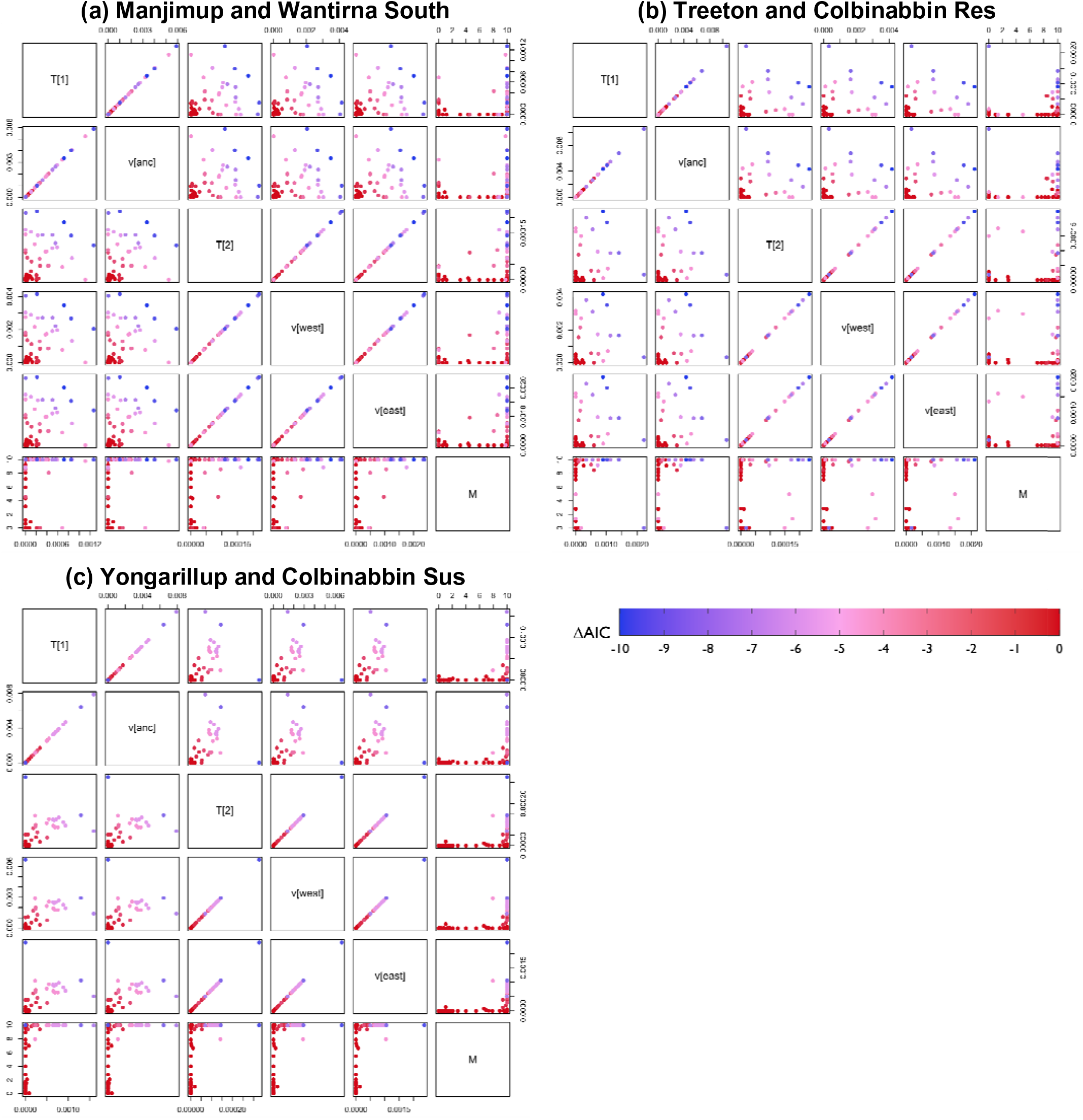
Correlation among parameter estimates for comparable replicate simulations (ΔAIC > −10) modelled under the symmetric migration scenario, for each population pair. Circles represent individual replicate simulations, coloured based on their ΔAIC to the best replicate simulation for their population pair (see legend).

**Figure S11.**
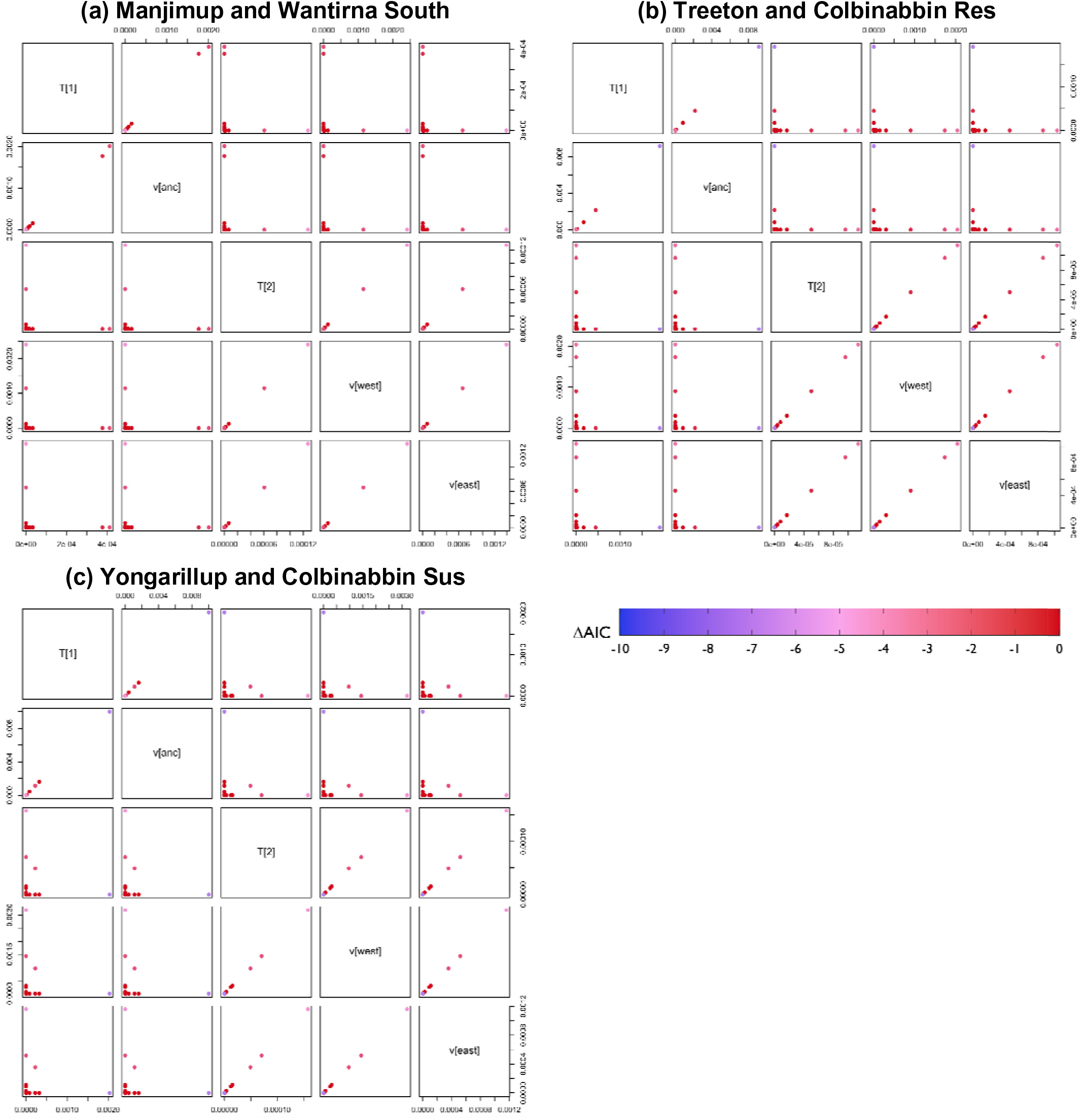
Correlation among parameter estimates for comparable replicate simulations (ΔAIC > −10) modelled under the isolation scenario, for each population pair. Points represent individual replicate simulations, coloured based on their ΔAIC to the best replicate simulation for their population pair (see legend).

**Table S1.** Samples used in population genomics analyses using whole-genome pool-seq of adult mites. Library IDs follow the nomenclature “RLEM[X]_[Y]_[A/B]” with “[X]_[Y]” denoting a unique DNA pool. “[A/B]” denotes a technical replicate (same DNA pool), either “A” or “B”. Libraries are grouped by biological sample and were merged for analysis.

| Sample ID | GenBank SRA | Population (Library ID; Replicate ID) | Year | Latitude | Longitude | Pooled mites |
| --- | --- | --- | --- | --- | --- | --- |
| RLEM2_2_A | SRR20755205 | Colbinabbin Sus (Lib=2; Rep=A) | 2021 | −36.5901 | 144.7994 | 50 |
| RLEM2_2_B | SRR20755204 | Colbinabbin Sus (Lib=2; Rep=B) | 2021 | −36.5901 | 144.7994 | 50 |
| RLEM4_1_A | SRR20755171 | Colbinabbin Res (Lib=4; Rep=A) | 2021 | −36.5642 | 144.8508 | 50 |
| RLEM4_1_B | SRR20755160 | Colbinabbin Res (Lib=4; Rep=B) | 2021 | −36.5642 | 144.8508 | 50 |
| RLEM18_X_A | SRR20755188 | Tambellup (Lib=18; Rep=A) | 2019 | −34.0456 | 117.5414 | 20 |
| RLEM18_X_B | SRR20755187 | Tambellup (Lib=18; Rep=B) | 2019 | −34.0456 | 117.5414 | 20 |
| RLEM19_1_A | SRR20755186 | Tambellup (Lib=19; Rep=A) | 2019 | −34.0456 | 117.5414 | 25 |
| RLEM19_1_B | SRR20755185 | Tambellup (Lib=19; Rep=B) | 2019 | −34.0456 | 117.5414 | 25 |
| RLEM20_1_A | SRR20755184 | Treeton (Lib=20; Rep=A) | 2019 | −33.851 | 115.2344 | 24 |
| RLEM20_1_B | SRR20755183 | Treeton (Lib=20; Rep=B) | 2019 | −33.851 | 115.2344 | 24 |
| RLEM21_1_A | SRR20755181 | Treeton (Lib=21; Rep=A) | 2019 | −33.851 | 115.2344 | 18 |
| RLEM21_1_B | SRR20755180 | Treeton (Lib=21; Rep=B) | 2019 | −33.851 | 115.2344 | 18 |
| RLEM26_2_A | SRR20755179 | Yongarillup (Lib=26; Rep=A) | 2019 | −33.7567 | 115.3869 | 14 |
| RLEM26_2_B | SRR20755178 | Yongarillup (Lib=26; Rep=B) | 2019 | −33.7567 | 115.3869 | 14 |
| RLEM27_2_A | SRR20755177 | Yongarillup (Lib=27; Rep=A) | 2019 | −33.7567 | 115.3869 | 29 |
| RLEM27_2_B | SRR20755176 | Yongarillup (Lib=27; Rep=B) | 2019 | −33.7567 | 115.3869 | 29 |
| RLEM28_2_A | SRR20755175 | Yongarillup (Lib=28; Rep=A) | 2019 | −33.7567 | 115.3869 | 25 |
| RLEM28_2_B | SRR20755174 | Yongarillup (Lib=28; Rep=B) | 2019 | −33.7567 | 115.3869 | 25 |
| RLEM29_2_A | SRR20755173 | Yongarillup (Lib=29; Rep=A) | 2019 | −33.7567 | 115.3869 | 14 |
| RLEM29_2_B | SRR20755172 | Yongarillup (Lib=29; Rep=B) | 2019 | −33.7567 | 115.3869 | 14 |
| RLEM34_1_A | SRR20755166 | Manjimup (Lib=34; Rep=A) | 2019 | −34.3101 | 116.1718 | 16 |
| RLEM34_1_B | SRR20755165 | Manjimup (Lib=34; Rep=B) | 2019 | −34.3101 | 116.1718 | 16 |
| RLEM35_X_A | SRR20755164 | Manjimup (Lib=35; Rep=A) | 2019 | −34.3101 | 116.1718 | 16 |
| RLEM35_X_B | SRR20755163 | Manjimup (Lib=35; Rep=B) | 2019 | −34.3101 | 116.1718 | 16 |
| RLEM36_1_A | SRR20755162 | Manjimup (Lib=36; Rep=A) | 2019 | −34.3101 | 116.1718 | 12 |
| RLEM36_1_B | SRR20755161 | Manjimup (Lib=36; Rep=B) | 2019 | −34.3101 | 116.1718 | 12 |
| RLEM37_X_A | SRR20755159 | Manjimup (Lib=37; Rep=A) | 2019 | −34.3101 | 116.1718 | 12 |
| RLEM37_X_B | SRR20755158 | Manjimup (Lib=37; Rep=B) | 2019 | −34.3101 | 116.1718 | 12 |

## Notes

### Competing Interest Statement

The authors have declared no competing interest.

### Summary of Updates

New analyses and restructured ms.

